# Intrinsic Molecular Timers and a Biphasic Amplitude Limit Regulate the Integrated Stress Response

**DOI:** 10.64898/2025.12.20.694968

**Authors:** Mustafa Ozen, Francesca Zappa, Karolina Kaminska, Daniel N. Itzhak, Stefka Tyanova, Carlos F. Lopez, Diego Acosta-Alvear

## Abstract

The Integrated Stress Response (ISR) is an evolutionarily conserved signaling network that remodels the translatome and transcriptome in response to multiple stresses, including nutrient deprivation, mitochondrial dysfunction, viral infection, and loss of protein homeostasis. Here, we present a comprehensive theoretical model of the ISR, calibrated to time-resolved proteomics data that captures how cells encode the magnitude and duration of stress signals to generate a homeostatic output. Our simulations and data converge on an ISR activation threshold defined by phosphorylated eIF2α levels, and sequential cascading delays in the accumulation of the ISR components ATF4, GADD34, CHOP, and DR5, suggesting hardwired molecular timers regulate ISR behaviors. Our combined experimental and computational analyses reveal limiting ATF4 levels, which can be suppressed when TC levels drop below a threshold that would allow its translation. While our model accurately predicts this initial saturation limit, its divergence from the data at high stress levels, correlated with minimal TC levels, identified a "translational cliff" that defines a finite ATF4-dependent ISR operational range. This work establishes a quantitative platform to probe ISR dynamics and generate novel, testable hypotheses.

## Introduction

The integrated stress response (ISR) is an evolutionarily conserved signaling network that endows cells with the means to cope with various stresses and restore homeostasis (Boone and Zappa, 2023). However, relentless, insurmountable stress causes the ISR to switch from adaptive to terminal, driving programmed cell death to eliminate cells that are injured beyond repair (Zappa et al., 2025). Given this key role in preserving and purging cells, it is not surprising that the ISR has been implicated in aging (Kourtis and Tavernarakis, 2011) and in major diseases of cellular dysfunction (Costa-Mattioli and Walter, 2020), including neurodegeneration (Wolozin and Ivanov, 2019) and cancer (Tian et al., 2021). Despite decades of research, causal links between ISR pathway dynamics (i.e., signaling amplitude, frequency, and duration) and pathophysiology remain obscure. For example, sectors of the ISR’s translational or transcriptional programs may play context- or cell-type-specific roles, and additional feedback layers may modulate ISR outputs depending on the cellular environment. Therefore, a detailed molecular understanding of such contextual ISR nuances is essential for effectively targeting the ISR for therapeutic intervention.

Four stress sensor kinases govern the mammalian ISR: general control nonderepressible 2 (GCN2), which detects amino acid shortage and ribosome collisions; heme-regulated inhibitor (HRI), which detects low heme, oxidative stress, and relays mitochondrial stress; PKR-like ER kinase (PERK), which detects disruption of proteostasis in the endoplasmic reticulum (ER), or “ER stress”; and protein kinase RNA-activated (PKR), which detects self and non-self double-stranded RNA (dsRNA) (Boone and Zappa, 2023). Upon detecting their cognate stress inputs, each kinase homodimerizes and autophosphorylates, which leads to their activation. The ISR kinases converge on phosphorylating a single serine (Ser51) on the α subunit (eIF2α) of eukaryotic translation initiation factor eIF2, a heterotrimeric GTPase, which, together with GTP and the initiator methionyl tRNA (Met-tRNAi), forms the eIF2-GTP-Met-tRNAi ternary complex (TC), which is essential for canonical translation initiation. Phosphorylated eIF2α (p-eIF2α) is a potent inhibitor of its guanine nucleotide exchange factor, the heterodecameric complex eIF2B (Clemens et al., 1982; Gordiyenko et al., 2019; Krishnamoorthy et al., 2001; Rowlands et al., 1988). Therefore, p-eIF2α accumulation decreases TC formation, resulting in less TC to deliver the Met-tRNAi to the ribosome, which leads to a temporary repression of protein synthesis.

The global translational repression induced by the ISR is coupled to the selective synthesis of specific proteins, including the transcription factors activating transcription factor 4 (ATF4) and C/EBP homologous protein (CHOP), through a well-studied mechanism involving upstream open reading frames hosted within the 5’ leader sequences of the corresponding mRNAs (Andreev et al., 2015; Harding et al., 2000; Hinnebusch et al., 2016; Lu et al., 2004). p-eIF2α also induces growth arrest and DNA damage-inducible 34 (GADD34), a regulatory subunit of protein phosphatase 1 (PP1), which dephosphorylates p-eIF2α, establishing a negative feedback loop that suppresses ISR signaling (Connor et al., 2001; Novoa et al., 2001). Severe or prolonged stress can cause irreparable damage to the cell, in which case the ISR engages a terminal signaling mode downstream of CHOP and contingent on the upregulation of death receptor 5 (DR5), a pro-apoptotic protein (Zappa et al., 2025). Through these mechanisms, the ISR reprograms the proteome and transcriptome, influencing cell states and determining cell fate (Costa-Mattioli and Walter, 2020; Holcik and Sonenberg, 2005; Hotamisligil and Davis, 2016; Pakos-Zebrucka et al., 2016).

The complexity of the ISR suggests intricate dynamic control. Experimentally addressing such dynamics in every set of parameters controlling nuanced ISR outputs is, in many cases, unfeasible. Mathematical models of cellular processes calibrated to experimental data offer an alternative tool to inform the formulation of hypotheses and predict pathway signaling outputs. These models are advantageous when collecting immense experimental datasets needed to support the interpretation of dynamic, multifaceted behaviors of signaling pathways is impractical or unattainable. Such computational models have offered valuable insights and guided testable predictions about the regulation of complex cellular processes, such as the mechanism of extrinsic cell death, the cross-talk between intra- and interneuronal networks, the effect of heterogeneity in cellular fates, and inhibition of ERK signaling in pancreatic cancer cells (Albeck et al., 2008; Emadi et al., 2022; Ozen et al., 2020; Sevrin et al., 2024; Shockley et al., 2019).

Here, we developed a computational model of the ISR, calibrated to time-resolved quantitative proteomics data, and validated some of its predictions experimentally. Together, our simulations and experiments support an ISR activation threshold contingent on the levels of p-eIF2α, a kinetic delay in the expression of the master ISR transcription factor ATF4, and intrinsic timers and limiters controlling ISR behaviors. These findings establish a mechanistic computational framework to test hypotheses in silico and guide future experimental design.

## Results

### A mathematical model to describe early and late ISR behaviors

To explore the ISR dynamics, we encoded the interactions between core ISR components from prior knowledge (Table 1). We deployed the Python-based modeling tool PySB (Lopez et al., 2013, see Methods) to build an ISR model, using a single ISR kinase, PKR, as a paradigm. The model considers ISR dynamics at baseline and upon PKR activation (Figure 1). At baseline, eIF2-GTP associates with Met-tRNAi to form the TC and deliver the initiator tRNA to the 40S ribosomal subunit to initiate protein synthesis. Upon recognition of the start codon, eIF2 hydrolyzes GTP, the 60S ribosome joins to form the elongation-competent 80S ribosome, and eIF2 is released to engage in another round of translation initiation (Brito Querido et. al., 2024). Starting another round of translation initiation requires GDP-GTP exchange by eIF2B on eIF2-GDP to restore the TC. Basal eIF2α phosphorylation and dephosphorylation by the gatekeeper holophosphatase CReP:PP1 is also included in the model (Jousse et al., 2003).

**Figure 1:**
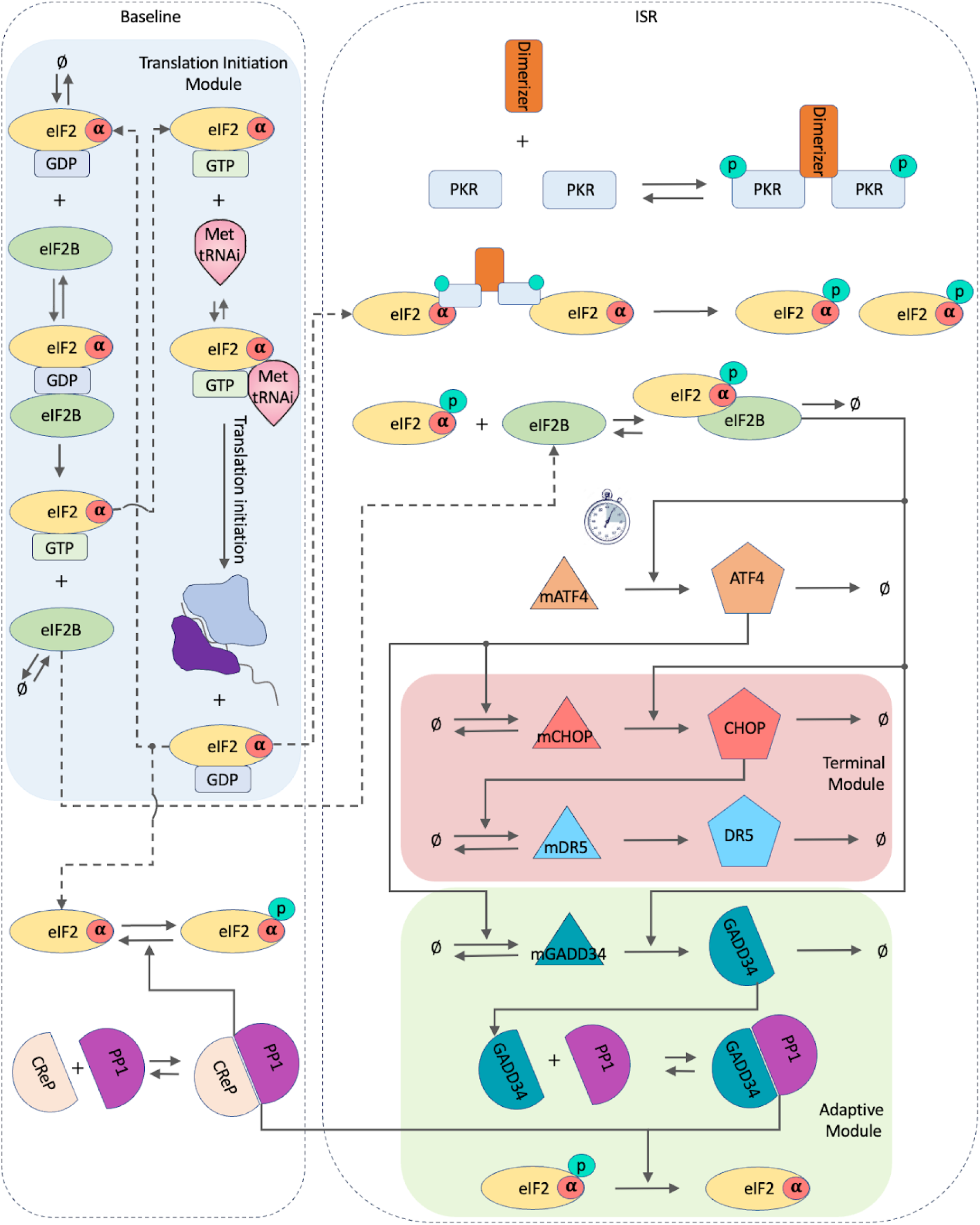
ISR model diagram. Model components at baseline and upon induction of the ISR. At baseline, the model considers GDP-GTP hydrolysis by eIF2B, TC formation, translation initiation, and basal eIF2α phosphorylation and dephosphorylation by CReP. Upon ISR induction, the model considers PKR activation, eIF2α phosphorylation, ATF4 expression, and induction of adaptive (ATF4-GADD34-dependent) and terminal (CHOP-DR5-dependent) modules. mRNAs are designated by the prefix “m” in the nodes depicted as triangles.

**Table 1:** ISR network wiring implemented in the model from prior knowledge.

| Interaction | Reference |
| --- | --- |
| eIF2, together with GTP and Met-tRNA<sub>i</sub>, forms the eIF2-GTP-Met-tRNA<sub>i</sub> ternary complex (TC). | Kapp and Lorsch, 2004 |
| TC initiates protein synthesis, where GTP is hydrolyzed | Kapp and Lorsch, 2004 |
| eIF2B binds to eIF2-GDP and exchanges GDP for GTP. | Gomez et al., 2002; Gordiyenko et al., 2014 |
| Basal p-eIF2α levels are present at steady state | Lu, Jousse et al., 2004 |
| PP1 binds to CReP to form its functional form (CReP:PP1). | Jousse et al., 2003 |
| CReP:PP1 dephosphorylates p-eIF2α. | Jousse et al., 2003 |
| Synthetic PKR dimerizes and activates with small-molecule ligand AP20187 (dimerizer). | Zappa et al., 2022 |
| Active PKR dimers phosphorylate two eIF2α (one per protomer). | Dar et al., 2005 |
| p-eIF2α sequesters eIF2B (p-eIF2α:eIF2B) and inhibits its activity. | Clemens et al., 1982; Gordiyenko et al., 2019; Krishnamoorthy et al., 2001; Rowlands et al., 1988 |
| p-eIF2α to total eIF2α ratio controls translation of ATF4. (Predicted to be around 30%). | Klein et al., 2022; This study. |
| Once the threshold is reached, *ATF4* is translated (with a rate depending on the amount of p-eIF2α:eIF2B, implemented in the model). | Andreev et al., 2015; Harding et al., 2000; Hinnebusch et al., 2016; Lu et al., 2004 |
| ATF4 upregulates GADD34 and CHOP | Averous et al., 2004; Ma and Hendershot, 2003 |
| CHOP induces DR5 | Lu et al., 2014; Yamaguchi and Wang, 2004 |
| PP1 binds to GADD34 to form its functional form (GADD34:PP1). | Connor et al., 2001 |
| GADD34:PP1 dephosphorylates p-eIF2α. | Rojas et al., 2015 |

We modeled ISR actuation upon PKR activation, which we used as a paradigm for several reasons. First, PKR’s biochemistry, structural biology, and activity in the cell are well-understood (Clemens, 1997; Cole, 2007; Dar et al., 2005; Feng et al., 1992; Li et al., 2006; Mayo et al., 2019; Nanduri et al., 1998). Second, PKR has been engineered into validated synthetic biology tools that allow activation with small molecules or light (Batjargal et al., 2023; Je et al., 2009; Jiang et al., 2010; Wong et al., 2025; Zappa et al., 2022). Third, uncoupling PKR activation from stress input sensing using these tools enables a clean dissection of the signal path, removing unwanted pleiotropic effects of natural PKR-activating inputs, i.e., self and non-self dsRNAs, or synthetic molecules resembling dsRNA, which can activate multiple cytosolic dsRNA sensors, such as RIG-I, MDA5, and OASs (Burkart et al., 2023; Donovan et al., 2015; Yu et al., 2018). For these reasons, we settled on using a validated chemical-genetics-based PKR activation approach consisting of PKR devoid of its dsRNA sensor domain and fused to a point mutant version of FKBP that selectively binds the small molecule ligand AP20187, leading to dimerization and activation (Zappa et al., 2022). As such, FKBP-PKR elicits a “pure ISR” that bypasses any cofunding signals that can encumber the model’s calibration and adversely impact the accuracy of its predictions.

Implementing the entire network of interactions between core ISR components (Figure 1) in a mechanistic model resulted in an extensive system of ordinary differential equations and several free parameters (Table S1). These free parameters are the unknown constants in our equations that represent the molecular properties and rates—such as reaction speeds and binding affinities—that define the system’s dynamic behavior and could not be directly measured experimentally. Accurately capturing the kinetics of all ISR downstream interactions demands calibrating these free parameters and requires precise measurements of on-pathway signaling activities, which we accomplished by collecting time-resolved (8 time points; 0, 1, 2, 4, 6, 8, 16, and 24 hours) proteomics-based measurements in H4 neuroglioma cells in which we activated FKBP-PKR.

Based on structural data, we reasoned that an active PKR dimer could, in principle, phosphorylate two eIF2α subunits, one per protomer (Dar et al., 2005), and incorporated this stoichiometry into the model. We modeled that reaching a set threshold level of p-eIF2α would enable the selective translation of the ATF4 mRNA and additional downstream canonical signals, including induction of the terminal (CHOP/DR5-dependent) and adaptive (ATF4-GADD34-dependent) ISR modules (Figure 1). Indeed, a p-eIF2α threshold level of 30 to 40% in cells treated with the ISR inducers thapsigargin and arsenite is sufficient to license stress granule formation, a classical marker of ISR induction (Klein et al., 2022). We encoded this threshold as a free parameter in our model and predicted a threshold value during model calibration using experimental measurements.

Our proteomics data indicated that FKBP-PKR activation indeed repressed global translation (the levels of more than 50% of the proteins measured were significantly decreased) and induced canonical downstream ISR signals: eIF2α phosphorylation coupled to selective expression of ATF4, CHOP, GADD34, and DR5 (Figure 2A and Zappa et al., 2022; Zappa et al., 2025). We corroborated ISR activity by analyzing the levels of canonical ISR signal path nodes, i.e., ATF4-GADD34 (the adaptive module) and CHOP-DR5 (the terminal module) (Figure 2B), in the proteomics dataset, which we then used to constrain the model’s free parameters (Methods, Zappa et al., 2022).

**Figure 2:**
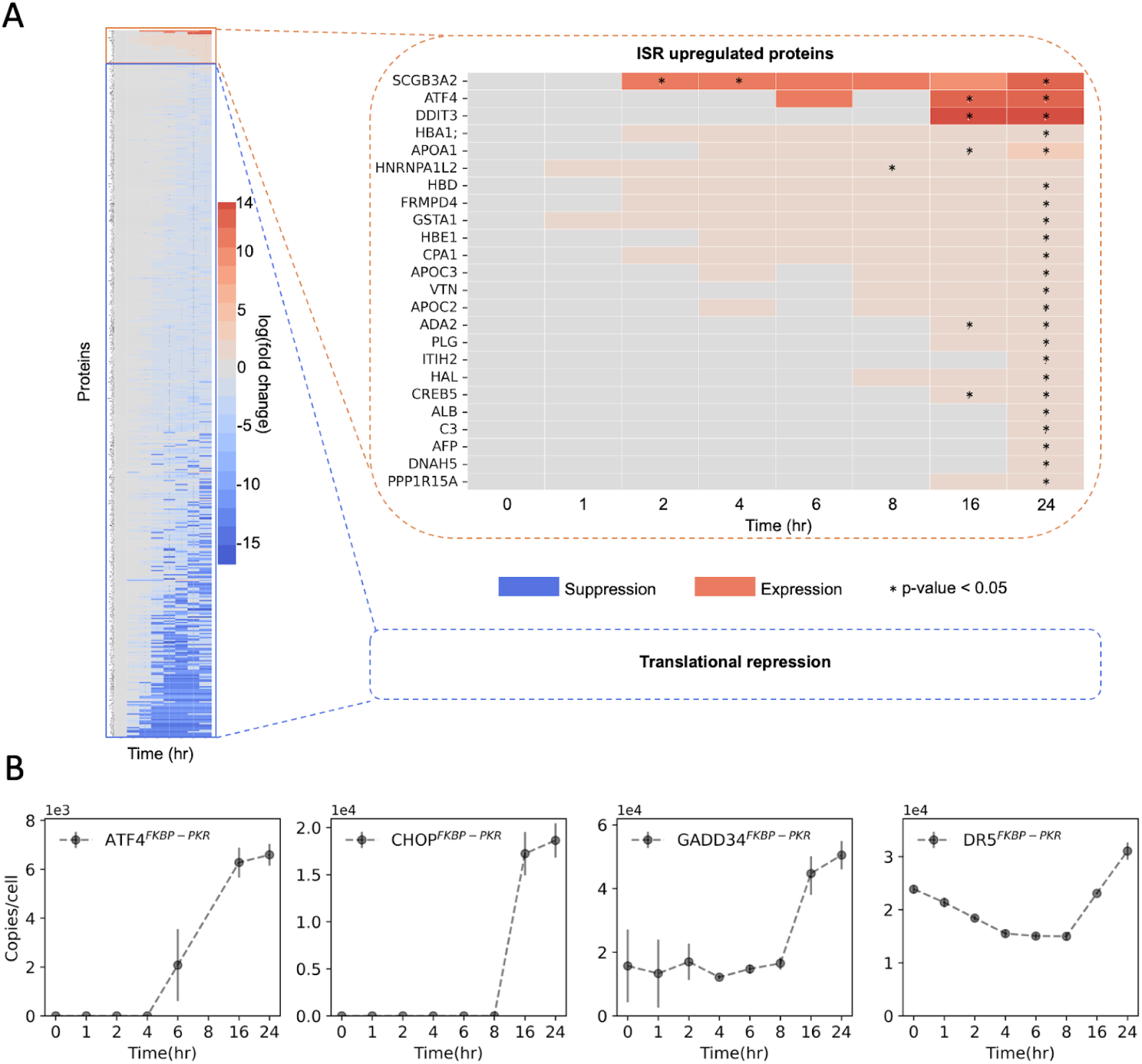
Synthetic ISR activation in H4 neuroglioma cells results in canonical outputs measured by mass spectrometry. (A) Heat map indicating that FKBP-PKR activation leads to a global translational repression and induction of a select few proteins, including the transcription factors ATF4 and CHOP. Rows: protein analytes. Columns: time points (0, 1, 2, 4, 6, 8, 16, and 24 hours) after FKBP-PKR activation. The plot illustrates differentially regulated proteins. (B) Protein expression dynamics of canonical ISR markers used to calibrate the model (n=3, Mean +/- StDev).

### The proportion of phosphorylated eIF2α licenses ISR activation

To accurately capture the initiation of the ISR and the kinetics of downstream signaling in our model, we encoded the p-eIF2α to total eIF2α ratio (defined as: p-eIF2α/(p-eIF2α + u-eIF2α; where u-eIF2α indicates unphosphorylated eIF2α) as the critical activation parameter; the phosphorylation of eIF2α by any of the ISR kinases is the initial signal transduction event in ISR actuation upon stress detection. For downstream efferent signaling, we settled on measuring ATF4 levels, as it captures both the ISR translational (i.e., reduction in TC levels is required to produce ATF4) and transcriptional (ATF4 drives the principal ISR gene expression program) outputs (Figure 2).

First, we estimated an activation threshold for the ISR in our model. To this end, we encoded the ratio of p-eIF2α to total eIF2α as a free parameter and evaluated the threshold by fitting the model to the experimental proteomics data. The high complexity of the ISR, which spans, in our model, translation initiation, eIF2α regulation, and the downstream bifurcating adaptive and terminal gene expression modules, yielded, expectedly, a large mathematical model composed of 56 free parameters. Unsurprisingly, this vast parameter space carried high dimensionality and the corresponding risk of parameter non-identifiability. Defining the p-eIF2α to total eIF2α ratio as a free parameter added a critical non-linearity to the system’s equations. This steep non-linear relationship dramatically increased the complexity of optimization, hindering the fitting algorithm from reliably finding parameter values, ultimately resulting in unsuccessful model calibration. Therefore, in a second step, we simplified the primary model by considering only the interactions and proteins whose induction is directly associated with ISR activation. Accordingly, we only included canonical and experimentally measurable proxies of ISR activity, specifically ATF4, GADD34, and CHOP mRNA and protein levels for model calibration (Figure S1A). We calibrated this simplified model through multiple iterations using different initial starting values (see Methods). This operation yielded 97 parameter sets, each representing a unique combination of all free parameter values that allowed the simplified model to successfully reproduce the experimental proteomics data equally well (Figure S1B). Using these parameter sets, we predicted that the p-eIF2α to total eIF2α ratio that would lead to experimentally measurable ISR canonical signals was, on average, 0.331 +/- 0.14 StDev (Figure 3A), which is consistent with previously reported observations in hepatocellular carcinoma cells stimulated with different ISR inducers (Klein et al., 2022).

**Figure 3:**
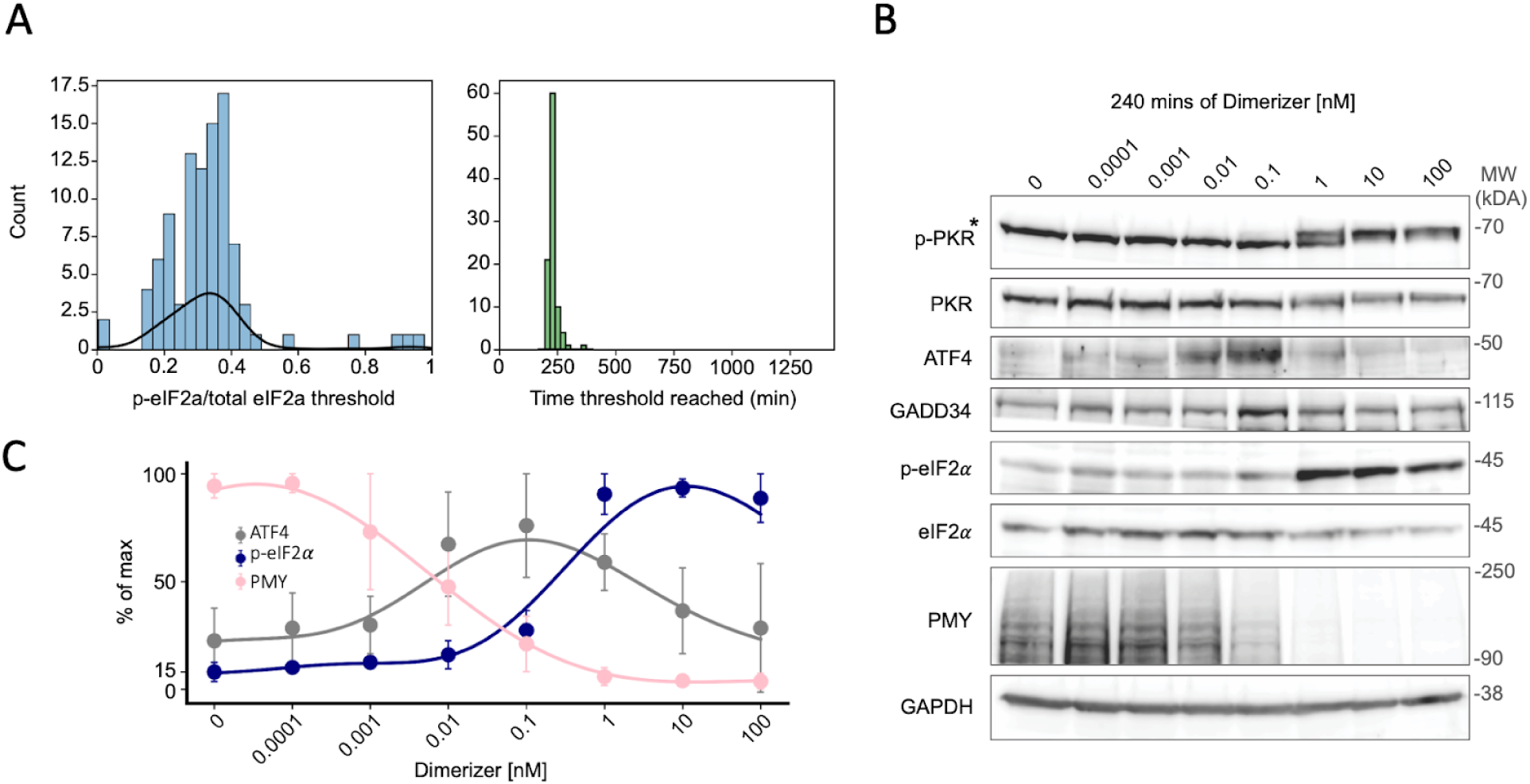
Experimental validation of the ISR model. (A) Model calibration resulted in 97 parameter sets that estimated a ratio of 0.331 p-eIF2α to total eIF2α leads to ISR activity (left panel). Y-axis (count): number of parameter sets within each bin. The black curve shows the estimated underlying distribution of the histogram (by a kernel density estimator). The model predicts that the 0.331 p-eIF2α to total eIF2α ratio is reached approximately 230 minutes after stress exposure (right panel). (B) Western blots showing ISR activation as a function of dimerizer dose (i.e., stress input level). The anti-puromycin (PMY) blot shows a ramping suppression of protein synthesis, evidenced by the decrease in the puromycin labeling of nascent peptides as a function of dimerizer doses. GAPDH: loading control. *non-specific band. (C) Quantification of the Western blot data in panel B. ATF4 protein levels are elevated and coincide with the approximate half-maximal protein synthesis suppression measured by puromycilation of nascent peptides when the p-eIF2α to total eIF2α ratio reaches approximately 20% (corresponding to a 0.01 nM concentration of dimerizer) and peak when the p-eIF2α to total eIF2α ratio reaches approximately 30% (corresponding to 0.1 nM dimerizer).

Next, we validated the model’s prediction by measuring ATF4 and GADD34 levels by Western blotting in FKBP-PKR H4 cells upon titration of the FKBP-PKR dimerizer, which allows ISR amplitude modulation and subsequent monitoring of downstream signals as a function of the p-eIF2 to total eIF2 ratio (Figure 3B). In these experiments, we observed that ATF4 expression rose above baseline when the p-eIF2α to total eIF2α ratio reached approximately 20% (corresponding to a 0.01 nM concentration of dimerizer), and peaked when the ratio reached approximately 30% (corresponding to a 0.1 nM concentration of dimerizer) (Figure 3C), which is consistent with our model’s prediction. We observed that increasing the stress input (i.e., dimerizer concentration) did not further increase ATF4 levels (Figure 3C). Instead, ATF4 protein levels drop when the dimerizer concentrations rise above 0.1 nM, presumably due to a strong suppression of translation initiation leading to a complete inhibition of protein synthesis at dimerizer levels above 1 nM, as measured by nascent peptide puromycilation (Figure 3B, PMY, lanes 6-8).

Based on these observations, we set the ISR activation threshold in the model to 30% p-eIF2α for further downstream analyses. While calibrating the primary model, we included dynamics for DR5 in addition to ATF4, GADD34, and CHOP, which accurately fitted the experimental data (Figure 4A). This data fit confirmed that the model faithfully captures the time-resolved protein accumulation of all four canonical ISR components measured—ATF4, GADD34, CHOP, and DR5—with the simulated trajectories closely conforming to the experimental proteomics data across the 24-hour time course. With this calibrated model, we next simulated the underlying mechanistic trajectories of all core pathway components (Figure 4B-I). Taken all simulations in consideration, our model predicted a rapid conversion of inactive PKR (u-PKR) to active p-PKR (Figure 4C), which induces a slow rise of the p-eIF2α to total eIF2α ratio until the threshold (0.3) is reached, at which point the slope abruptly changes, reflecting a steep (slope = 0.7) transition that eventually reaches saturation (Figure 4D). Mechanistically, we interpret this sudden change as the product of a lag time in transcriptional induction of ATF4, GADD34, CHOP, and DR5 mRNAs, which drives the continuous accumulation of their corresponding proteins over the 24-hour period (Figure 4F-I). By the time the p-eIF2α to total eIF2α ratio reaches the saturation point, we observed that levels of free eIF2B drop to near completion (Figure 4E). These changes are reflected in TC availability, with an anticipated abrupt drop in TC levels upon ISR activation beginning at approximately 2 hours into the simulation, and leading to profound global translation repression (Figure 4B).

**Figure 4:**
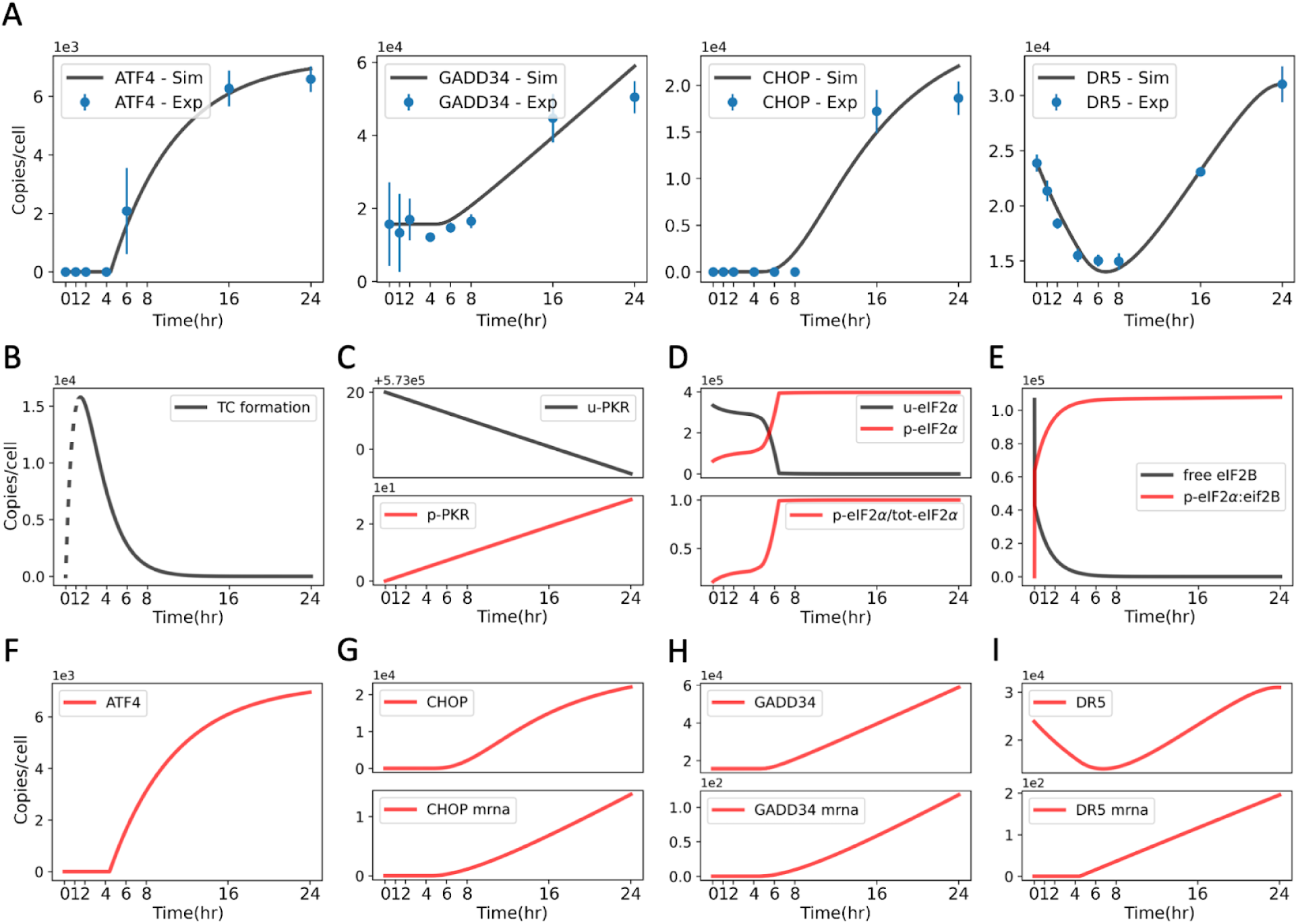
Model calibration to experimental data with a fixed ratio of 30% of p-eIF2α to total eIF2α. (A) Model goodness of fit to the experimental data (MSE = 0.053). Setting the activation threshold to 30% p-eIF2α to total eIF2α in the primary model and calibrating it to the experimental data resulted in an optimal parameter set. Reaching the activation threshold of 30% p-eIF2α to total eIF2α resulted in a logarithmic increase in ATF4 levels (left), near linear rise in GADD34 (second from left), and sigmoidal increases in CHOP and DR5 levels (second from right and the rightmost panels, respectively), all fitted by considering Hill functions with different variables and coefficients (Table S1). (B-I) Trajectories of model components after parameter calibration. (B) TC levels as a function of stress duration. The dashed line indicates the equilibration of the model in the early time points with an imputed TC value of 0. This equilibration does not reflect the TC levels in the cell during the 0-2 hour window; however, it does not affect the rest of the model dynamics portraying the drop in TC levels following ISR induction starting at 2 hours and throughout the rest of the simulation. (C) u-PKR and p-PKR trajectories as a function of stress duration. (D) u-eIF2α, p-eIF2α levels (top), and p-eIF2α to total eIF2α ratio (bottom) as a function of stress duration. (E) Free (uninhibited) vs p-eIF2α-inhibited eIF2B as a function of stress duration. (F) ATF4 levels as a function of stress duration. (G-I) CHOP, GADD34, and DR5 protein levels (top) and their respective mRNA levels (bottom) as a function of stress duration. “Copies/cell” in the y-axis indicates the absolute molecular count (protein or mRNA species) per cell (see Methods).

### Model interrogation defines the boundaries of activation and resolution dynamics

Having constrained the model’s activation kinetics, we next sought to understand the temporal regulation of committed ISR engagement. We performed in silico experiments simulating different stress durations and found that an exposure to stress of approximately 4 hours and 20 minutes was sufficient to induce a persistent ATF4 response that, in the model, did not resolve upon stimulus removal (Figure 5). This prediction aligns with our proteomics data, indicating a lag time of at least 4 hours preceding the accumulation of ATF4 protein (Figure 2B). However, the persistent activity, despite the removal of the stimulus, antagonizes the sub-one-hour half-life of the ATF4 protein (Lassot et al., 2001). We attribute this discrepancy to our experimental design, which employed a sustained high-level input to induce the ISR, potentially leading to the dominance of the terminal module over the adaptive module. The model was calibrated accordingly to this setting. Regardless, this approach provided robust constraints for the ISR’s activation machinery, even though it bypassed information about the GADD34-mediated feedback that governs the ISR deactivation. The model’s inability to resolve these aspects of signaling highlights its boundaries while confirming that ISR resolution is a distinct kinetic module that would require different experimental paradigms, such as pulse-chase studies, to parameterize.

**Figure 5:**
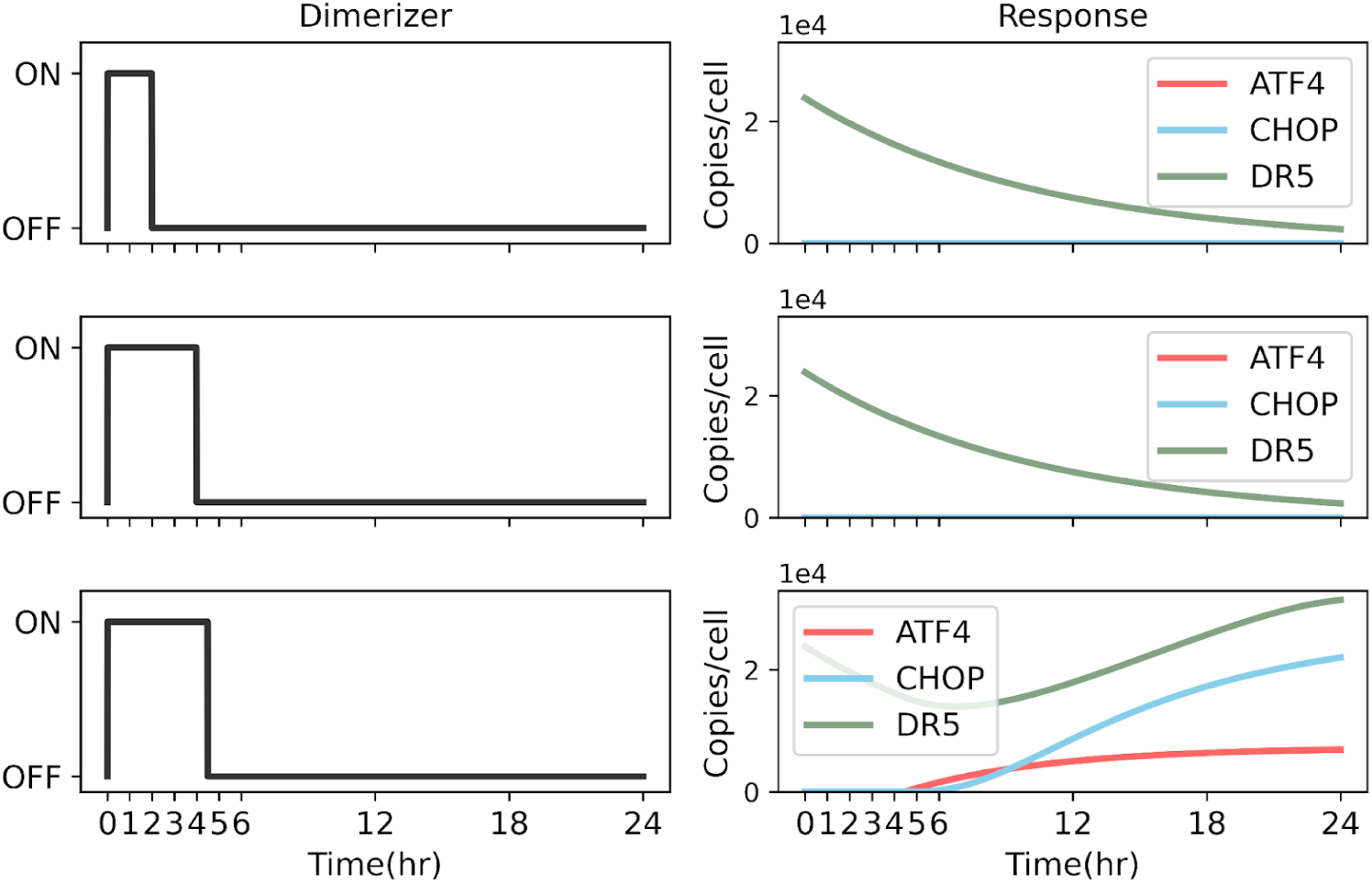
Stress duration tailors the ISR amplitude. Simulated trajectories for ATF4, CHOP, and DR5 protein levels when cells are exposed to dimerizer for different durations (3, 4, or 4 hours and 20 minutes) followed by a refractory period. The model predicts that sustained stress exposure of approximately 4 hours is required to engage a persistent ISR program that continues even when the stimulus is removed.

Next, we tested how increasing the magnitude of the stress input modulates the response. To this end, we ran a series of simulations in which we changed the amount of dimerizer applied for 24 hours, from 0 to 2000 copies, in increments of 100 copies. To normalize our input scale to the dose used in the proteomics experiment, we assigned a random value of 1000 copies to correspond with a 100 nM dimerizer. This arbitrary value assignment provides a convenient reference point for the simulation range without affecting the simulation outcome, as the associated kinetic rate constants scale accordingly during model calibration. In these simulations, we did not observe a positive correlation between the ISR signal amplitude (measured by ATF4 levels) and increasing dimerizer doses (Figure 6A).

**Figure 6:**
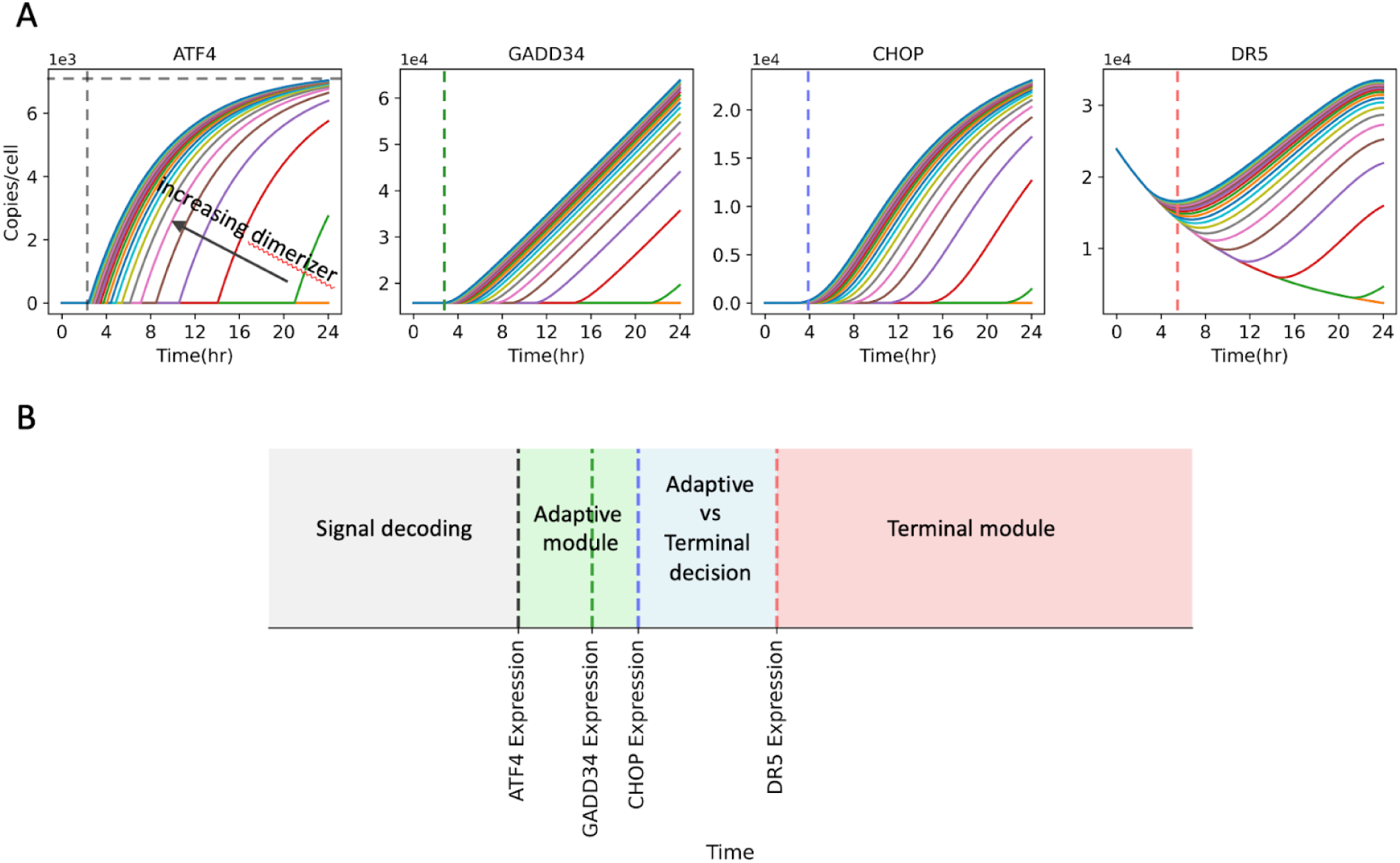
Simulation of the effect of stress amplitude on the adaptive and terminal ISR modules. (A) Predicted protein levels of the adaptive (ATF4-GADD34) and terminal (CHOP-DR5) nodes as a function of stress duration. ISR onset, defined by the time at which ATF4 protein expression starts, occurs at approximately 21 hours with the minimum dimerizer concentration simulated (100 copies, green trace). Increasing the dimerizer concentration, i.e., increasing the simulated stress level, shifts the ISR onset to an earlier time, with the fastest response obtained at approximately 2.7 hours (indicated by the vertical black dashed line in the leftmost panel), and with a simulated dimerizer concentration of 2000 copies. The horizontal dashed black line indicates a response saturation of about 7000 copies of ATF4 molecule per cell. GADD34, CHOP, and DR5 exhibit a similar behavior; however, their minimum expression time at the maximum dimerizer concentration is offset from that of ATF4 by +0.8 hours for GADD34 (to 3.5 hours, indicated by the dashed green line), +1.3 hours for CHOP (to 4 hours, indicated by the dashed blue line), and +2.8 hours for DR5 (to 5.5 hours, indicated by the dashed red line). (B) Model of the ISR’s cascade of events predicted by our simulation. The model predicts that the ISR unfolds through a series of hardwired molecular timers. Following an initial signal decoding lag, the program proceeds through sequential activation of the adaptive module (ATF4/GADD34), a decision-making phase (CHOP), and the terminal module (DR5). Note that the respective lag times separating these events might be cell-type or stressor-specific.

While the model correctly identified an amplitude limiter that is consistent with our experimental data showing that ATF4 levels saturate at 0.1 nM dimerizer (Figure 3C), the latter revealed a biphasic behavior wherein ATF4 levels drop after the saturation point when the dimerizer concentration is increased beyond 1 nM (Figure 3C). This phenomenon, which our model does not capture, suggests that overwhelming stress leads to a massive drop in TC levels and profound translational suppression, even the ATF4 mRNA cannot escape. These observations indicate that the ISR’s adaptive module has a finite operational range.

Within this range, however, the model accurately predicts the system’s hardwired molecular timers. Our simulations show an intrinsic signal decoding lag time of approximately 3 hours before ATF4 induction, regardless of stress amplitude. This initial timer governs a cascade of subsequent events, with GADD34, CHOP, and DR5 expression occurring sequentially at approximately 3.5, 4, and 5.5 hours, respectively (Figure 6A). The concatenation of these events defines the temporal stages of the ISR (Figure 6B), from signal decoding to the adaptive response, a critical decision-making phase, and finally, the terminal program.

### u-eIF2α and eIF2B levels determine the ISR amplitude

To assess how changes in the protein levels of ISR components affect ISR outputs, which can be a primary contributor to stress response heterogeneity in a cell population, we conducted Global Sensitivity Analysis (GSA). This analysis systematically queries the entire multi-dimensional parameter space to determine which model inputs (i.e., component concentrations and kinetic rate parameters) drive the most substantial variance in the model outputs. A model input driving high variance indicates that the model is highly sensitive to it. To this end, we computed the first-order (S1), second-order (S2), and the total effect (ST) sensitivity indices using the Sobol method (Sobol, 2001, see Methods). The S1 index quantifies the direct impact of a single input parameter; the S2 index quantifies the influence of interactions between two parameters; and the ST index quantifies the combined effect of a single parameter and all its potential interactions with every other parameter in the model output.

First, we analyzed the sensitivity of the model’s adaptive and terminal ISR modules, with ATF4 and GADD34, representing the adaptive module and feedback control, and CHOP and DR5 representing the programmed cell death module, to the initial levels of individual mRNAs and proteins (Figure 7). In these analyses, we found a clear-cut separation in control mechanisms: the expression of GADD34, CHOP, and DR5 was primarily sensitive to the initial levels of their respective mRNAs and proteins, but largely insensitive to upstream regulators, i.e., the dimerizer, p-eIF2α, and ATF4 (Figure 7, note the parameters with the maximum S1 and ST values in the heatmaps occur in GADD34, CHOP, and DR5). In contrast, the expression of ATF4 was uniquely and highly sensitive to its upstream regulatory components (Figure 7). The highest sensitivity indices for ATF4 were observed for u-eIF2α, eIF2B, and CReP, with the highest ST and S2 indices for u-eIF2α and eIF2B (Figure 7 and supplementary spreadsheets), indicating that the relative abundance and interaction of u-eIF2α and eIF2B are the principal determinants controlling ATF4 levels. Similarly, the sensitivity of ATF4 to CReP levels is congruent with the established control of translation initiation through regulation of basal p-eIF2α levels, as has been reported (Xu et al., 2018).

**Figure 7:**
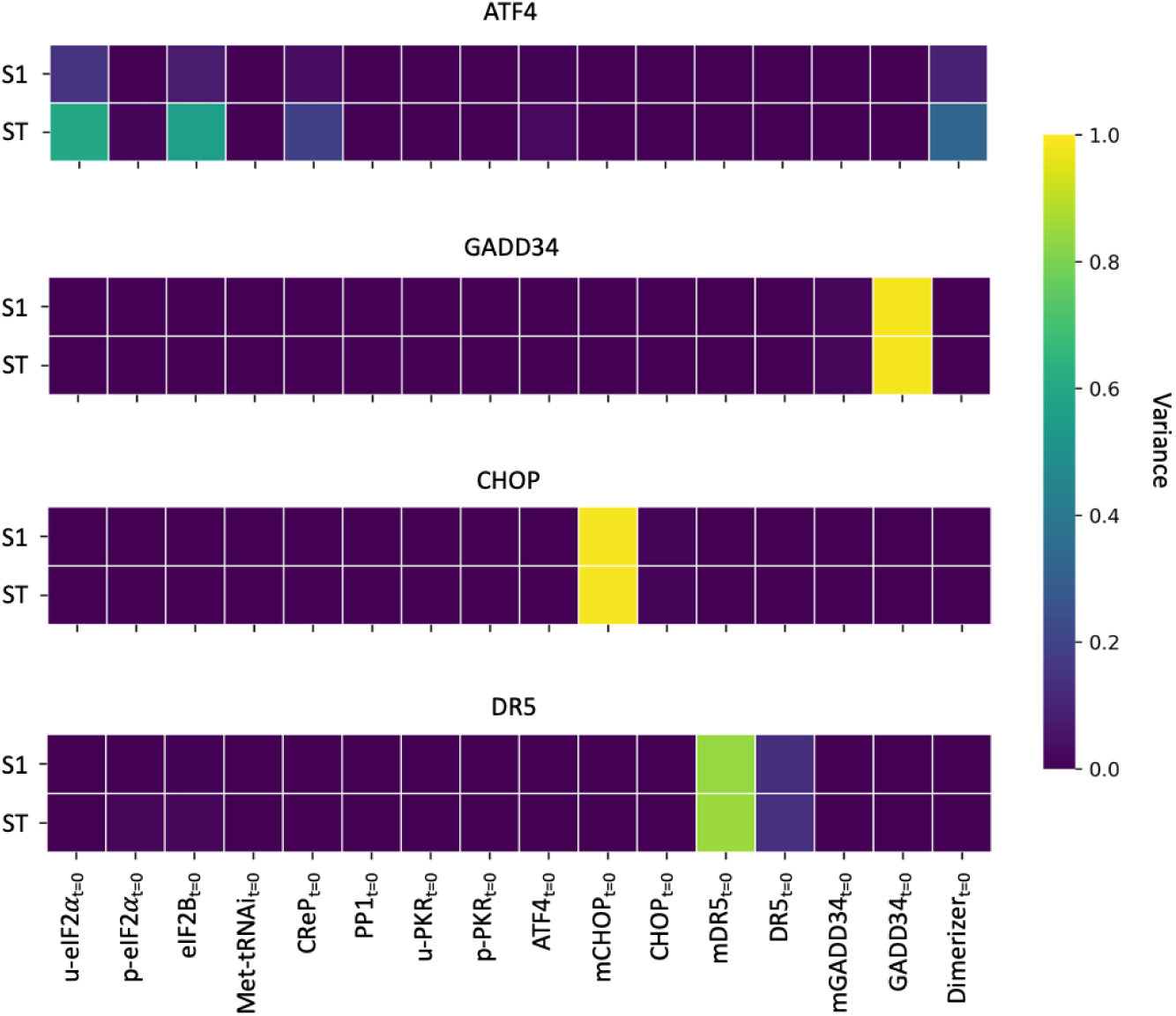
Model sensitivity analysis based on its initial values. The model’s sensitivity, estimated using the initial values of each critical node (protein or mRNA values at t=0-hr), was computed for ATF4, CHOP, GADD34, and DR5 independently. In these calculations, these four proteins represent model outputs, while the initial values of all nodes in the model (see Figure 1) were considered inputs. The sensitivity values were computed based on the goodness of fit of the experimental data to the model’s output (i.e., variance in the MSE between output and the experimental data, as indicated by the colorbar). A high variance indicates that the model is highly sensitive to a particular input. S1 measures the contribution to the output variance of the change in an individual initial value. ST measures the contribution to the output variance caused by an individual initial value, including both its first-order effects (variance of the initial value) and all higher-order interactions (covariance of the initial value alongside combinations of other initial values).

We observed a similar pattern for model sensitivity dictated by the free parameters for ATF4, CHOP, GADD34, and DR5. For these four ISR components, mRNA expression rate constants notwithstanding, the model was globally sensitive to PKR phosphorylation and eIF2α phosphorylation and dephosphorylation rate constants (Figure S2). These results indicate, expectedly, that the overall amplitude and kinetics of the ISR are primarily dictated by the rates at which the initial ISR signal is generated (i.e., the rate of eIF2α phosphorylation) and at which that signal subsides (i.e., the rate of p-eIF2α dephosphorylation). Interestingly, the model was more sensitive to the free parameters associated with p-eIF2α dephosphorylation by CReP compared to dephosphorylation by GADD34, emphasizing the importance of CReP’s role in regulating basal p-eIF2α levels, which we interpret as CReP being the gatekeeper for setting the sensitivity threshold for the ISR. This interpretation is consistent with observations that loss-of-function mutations in the gene encoding CReP (PPP1R15B) are associated with sustained elevation of basal p-eIF2α and chronic ISR signaling (Torkenczy et al., 2025), as observed in a human syndrome of microcephaly, intellectual disability, short stature, and diabetes (Abdulkarim et al., 2015). However, the contribution of GADD34-mediated negative feedback might be underestimated in our simulations since, in our experimental system, the dimerizer was not removed for the duration of the experiment, leading to sustained activation of FKBP-PKR, which likely overrides GADD34’s negative feedback and favors the terminal response. Indeed, in cells in which FKBP-PKR is constitutively active, the ISR leads to apoptosis (Zappa et al., 2025).

## Discussion

The ISR allows cells to adapt to various stressors by dynamically modulating their biosynthetic capacity through translation control and induction of gene expression programs. Here, we used a comprehensive theoretical model of the ISR, calibrated to time-resolved proteomics data, to interrogate the system’s dynamic behaviors. Using ATF4 expression as a proxy for both the translational and transcriptional arms of the response, our analyses revealed that ISR activation is not merely proportional to the magnitude of the stress input, but is also regulated by hardwired kinetic properties and a limited response range. Specifically, our model predicted an activation threshold that acts as an ON/OFF switch dependent on the levels of p-eIF2α, a protracted response time that allows deciphering transient signals to evoke adaptive versus terminal responses, and an embedded amplitude limiter that prevents a runaway response.

Our model’s prediction and experimental validation that approximately 30% (+/- 10%) of p-eIF2α is required to initiate ATF4-dependent ISR signals (Figure 3) indicates the existence of the aforementioned ON-state switch, which is consistent with previous findings (Klein et al., 2022). Our simulations also indicate that the ATF4-dependent ISR signal output is kinetically buffered, requiring approximately 4 hours of stress exposure to induce ATF4 expression (Figure 5). This lag suggests that stress management proceeds through an early stress resolution program, likely contingent on fine-tuning translation initiation, which precedes gene expression programs to homeostasis. These observations support a model in which hardwired molecular timers govern the ISR (Figure 6B). While lag times might differ for various stressors and cell types, we surmise that a cell first engages a preemptive stress resolution mode contingent on the continued phosphorylation of eIF2α by active ISR kinases. This built-in timer enables stress signal decoding and provides latency for translating sufficient ATF4 mRNA molecules to initiate a transcriptional response. Transcriptional regulation allows the ISR to match its output to cellular demands in two successive stages: an early adaptive stage wherein ATF4 drives homeostatic gene expression programs, which transitions into a second decision-making phase marked by CHOP expression. If the stress is no longer resolvable and the cell is irreparable, then CHOP drives a pro-apoptotic program contingent upon DR5 expression (Zappa et al., 2025). These events mark the ISR’s final transition, from adaptive to terminal.

A key finding of our study is the discrepancy between the model and the experimental data on the ISR’s input-output relationship. While our model, based on the canonical pathway structure, correctly predicted that the ATF4 response would saturate at high stress levels, our experimental data revealed that the trajectory of ATF4 protein levels diverged from this prediction. Even though ATF4 protein levels saturated at moderate stress (0.1 nM dimerizer), they *decline* upon increasing stress inputs (1-100 nM), indicating that saturating levels of p-eIF2α severely limit TC formation and halt all cap-dependent translation, including that of the ATF4 mRNA (Figure 3B). We interpret these results as an inherent ISR "translational cliff" which cannot be overridden by the GADD34-dependent negative feedback. Such a point of no return indicates that the ISR possesses a finite operational range, providing a potential mechanistic explanation for how a cell might distinguish between manageable stresses and a catastrophic one.

Heterogeneity in a cell population contributes to biological noise (Eling et al., 2019), which is inherent to all signal transduction pathways, including stress responses (Adamson et al., 2016; Guilbert et al., 2020). Thus, it is plausible that cell-to-cell heterogeneity in ISR signaling arises from individual cell variability (i.e., divergent stress response amplitudes, lag times, and the differential engagement of parallel signaling pathways). In our simulations, we attempted to predict the molecular source of this variability through global sensitivity analysis for ATF4, GADD34, CHOP, and DR5 with respect to all of the ISR components embedded in the model. Our results indicate that the initial abundance of u-eIF2α and eIF2B, and the interplay between them, are the main determinants controlling the amplitude of the ATF4-dependent ISR, as they cause the highest variance in ATF4 levels (Figure 7 and supplementary spreadsheet for S2 index). This finding provides a putative molecular explanation for cell-to-cell heterogeneity in the ISR (Klein et al., 2022). If this is the case, the cell’s intrinsic capacity for adaptive signaling is inherent to the activity of ISR kinases coupled to the intrinsic stoichiometric ratio of the core translation machinery that licenses the downstream ATF4 program.

Given that some of our model’s predictions have been validated experimentally and align with what has been observed in the literature, our model is sufficiently robust to serve as a reliable platform for future in silico hypothesis testing and guide experimental design. For instance, the model could be deployed to test how reducing the initial concentration of eIF2B (by 50% in a simulated cell population) alters the ISR dynamics, or to explore whether abolishing CReP activity (which removes basal p-eIF2α dephosphorylation) impacts the ATF4 kinetic buffer. The model can be also be used to simulate regimes of activation/deactivation or inhibition of specific components by introducing new species or by simulating the effect of ISR-modulating drugs such as the eIF2B activators ISRIB (Sidrauski et al., 2015) and 2BAct (Wong et al., 2019), or small molecules that directly impact ISR kinase activity, such as neratinib, which activates GCN2 (Tang et al., 2022) and the PERK inhibitors GSK2656257 and GSK2606414 (Szaruga et al., 2023), allowing for the systematic investigation of emergent regulatory effects and the prediction of novel control points within the ISR signaling cascade. Another potential use of our model includes cell fate determination upon stress encounters. The model can predict the quantitative conditions under which the terminal ISR is engaged by identifying the optimal timing and magnitude of an ISR activation pulse that selectively induces the adaptive (ATF4-GADD34) module without engaging the terminal, DR5-dependent apoptotic program. Such simulations would allow predicting conditions that favor cellular adaptability.

Our work reinforces the notion that the ISR is not a simple linear pathway, but a system of sophisticated decision-making molecular circuits that use thresholds, timers, and limiters to measure and interpret the nature of the stress experienced by the cell. It is likely that these properties have been selected to optimize the cell’s use of limited resources to ensure adaptability. Understanding this control logic has profound implications, since the ISR is dysregulated in neurodegeneration, metabolic disease, and cancer. Our model provides a quantitative framework to define the kinetic parameters that regulate the ISR’s adaptive and terminal phases, with the potential to illuminate how this balance is lost in disease. Such knowledge is essential for designing therapeutic strategies for precise interventions modulating ISR activities to restore homeostasis, regain adaptability, and reverse disease.

### Limitations of the study

Our current model has limitations that narrow its scope and impact its predictive capacity. While our chemical-genetic approach successfully isolated the PKR branch to study a "pure" ISR signal, the model does not include the three other ISR kinases (PERK, GCN2, and HRI). Consequently, it cannot predict complex crosstalk dynamics or differential responses when the cell encounters more than one stress (e.g., ER stress and nutrient deprivation), which limits its application in understanding the full complexity of the ISR in physiological contexts. However, as data becomes available, additional ISR nodes could be embedded in the model, allowing for an understanding of response dynamics when one or multiple ISR kinases are simultaneously active. Moreover, the model relies on simplified rate laws (e.g., mass action kinetics) to illustrate complex multi-step processes like enzyme catalysis. As a result, the fitted parameters represent effective phenomenological rates rather than actual physical rate constants, which may limit the model’s predictive accuracy when simulating concentrations far outside the calibrated regime. Additionally, our model only considers the ISR’s temporal dynamics and not its spatial organization. Therefore, while useful, our model remains approximate, as many downstream ISR processes are not only temporally but also spatially regulated, as occurs with the formation of stress granules and the localized fine-tuning of translation near organelles (Brar et al., 2024; Chakrabarty et al., 2024; Sidrauski et al., 2015).

The quantitative reliability of the model’s parameters also presents a technical challenge. Despite our rigorous model calibration efforts, we only obtained a limited number of distinct parameter sets that fit the experimental data equally well, and only a single parameter set produced downstream trajectories that were biologically reasonable (Figure 4B-I). Thus, assessing the model’s mechanistic implications would require obtaining the complete probability distributions for each one of the free parameters adjusted to the full range of values consistent with experimental data. The model’s inherent complexity and multiparameter nature limit its calibration, even through advanced methods like Bayesian parameter inference. More sophisticated approaches are required to address the complexities of high-dimensional, nonlinear problems (Linden et al., 2022). Resolving parameter ambiguity can provide insights into the pathway’s biochemistry and open up additional ways to analyze and interpret the model, for example, by exploring parameter identifiability and signal execution modes (Ortega et al., 2024). Further model refinements are resource-intensive and will inevitably require additional experimental data, including parallel experiments in the absence of input to capture the baseline dynamics, and in the presence of multiple doses of input to capture the nonlinear effects, which lie beyond the scope of this work. Such experiments could be addressed in subsequent investigations.

## Methods

### Data collection and processing

#### a) Proteomics

##### i) Sample preparation

H4 cells were grown in 6-well plates. The day after, cells were incubated with 100nM of AP20187 (B/B homodimerizer, Takara HY-13992) for the indicated times. At the end of the time course, cells were harvested as follows: media was aspirated, followed by two washes with PBS, and cells were scraped in lysis buffer (100 mM Tris buffer pH 8.5, containing 1 % Sodium Deoxycholate), then heated to 95 °C for 5 minutes. Lysates were reduced with 1 mM DTT for 15 minutes at 25 °C, and alkylated with 5 mM Iodoacetamide for 20 minutes at 25°C in the dark. Samples were digested with LysC at a 1:50 enzyme:protein ratio overnight at 37 °C, and digestion was continued for a further 4 hours following the addition of Trypsin at a 1:50 enzyme:protein ratio. Digested material was subjected to solid phase extraction as follows: digests were mixed with an equal volume of isopropyl alcohol, acidified to 1% TFA final, and loaded directly onto 50 mg cartridges of Strata-XC (Phenomenex), washed once with 99.9% IPA, 0.1% TFA, once with 0.1% TFA, and once with 0.1% formic acid. Peptides were eluted with 80% acetonitrile, 5% Ammonium Hydroxide, and immediately subjected to vacuum centrifugation. Peptide samples were resuspended in 3% Acetonitrile, 0.1% formic acid prior to MS analysis.

##### ii) Mass Spectrometry

Mobile phases A (0.1 % formic acid) and B (80% acetonitrile, 0.1% formic acid) were used to separate samples on a linear gradient from 3 - 35 % B over 60 min, on a 1mm x 150mm Waters CSH C18 column maintained at 55 °C. Ions were generated using an HESI source, spray voltage was set to 3500 V, the ion transfer tube was set to 325 °C, and the vaporizer was set to 125 °C. A Thermo Fisher Scientific Orbitrap Ascend was used to measure peptides with a data-independent acquisition scheme. MS1 scans were acquired from 350-1050 m/z on an Orbitrap analyzer at a resolution of 120,000. Following quadrupole isolation with a 10.25 m/z window and HCD activation at 31 % normalized collision energy, MS2 scans were measured in the Orbitrap at 15,000 resolution from 200-1800 m/z.

##### iii) Copy number calculations

All .RAW files were processed in DIA-NN v1.8 against a human database from Uniprot downloaded on March 10th, 2023, and a common contaminants .fasta file from MaxQuant. Match between runs was enabled.

Absolute protein quantification in the form of copy numbers was calculated from DIA intensities as calculated by DIANN using the Proteomic Ruler approach (Wiśniewski et al., 2014). In short, the summed intensity of histones in a deep proteome is proportional to the amount of DNA, which depends on the number of cells. The proportion of summed histone intensity to total mass spectrometry (MS) signal corresponds to the proportion of cell DNA mass to cell protein mass. Relating the histone MS signal to the total MS signal allows one to estimate the protein mass per cell at a given cell ploidy and genome size (TotalProteinAmount).

Diploid cells have 6.5pg DNA mass. H4 cells have 73 chromosomes, so the DNA mass in these cells was calculated as 6.5452025pg * (73/46).

The mass per cell for each protein is estimated as the product of its MS signal fraction and the cellular protein mass (TotalProteinAmount). To calculate copy numbers per protein, we then used the following formula:

CN = ((protein MS / totalMS) * TotalProteinAmount) / 10^12 * (Avogadro’s number/protein molecular weight).

Protein molecular weight was taken from UniProt (UniProt Consortium, 2025), and the list of histones was taken from Wisniewski et al. ( 2014).

#### b) Immunoblotting

Cell lysates were collected directly in Laemmli SDS-PAGE sample buffer (62.5 mM Tris-HCl pH 6.8, 2% SDS, 10% glycerol, and 0.01% bromophenol blue). Lysates were briefly sonicated and supplemented with fresh 5% 2-mercaptoethanol before heat denaturation and separation by SDS-PAGE. Lysates were separated on SDS-PAGE gels and transferred onto nitrocellulose membranes for immunoblotting. Immunoreactive bands were detected by enhanced chemiluminescence using horse radish peroxidase (HRP)-conjugated secondary antibodies.

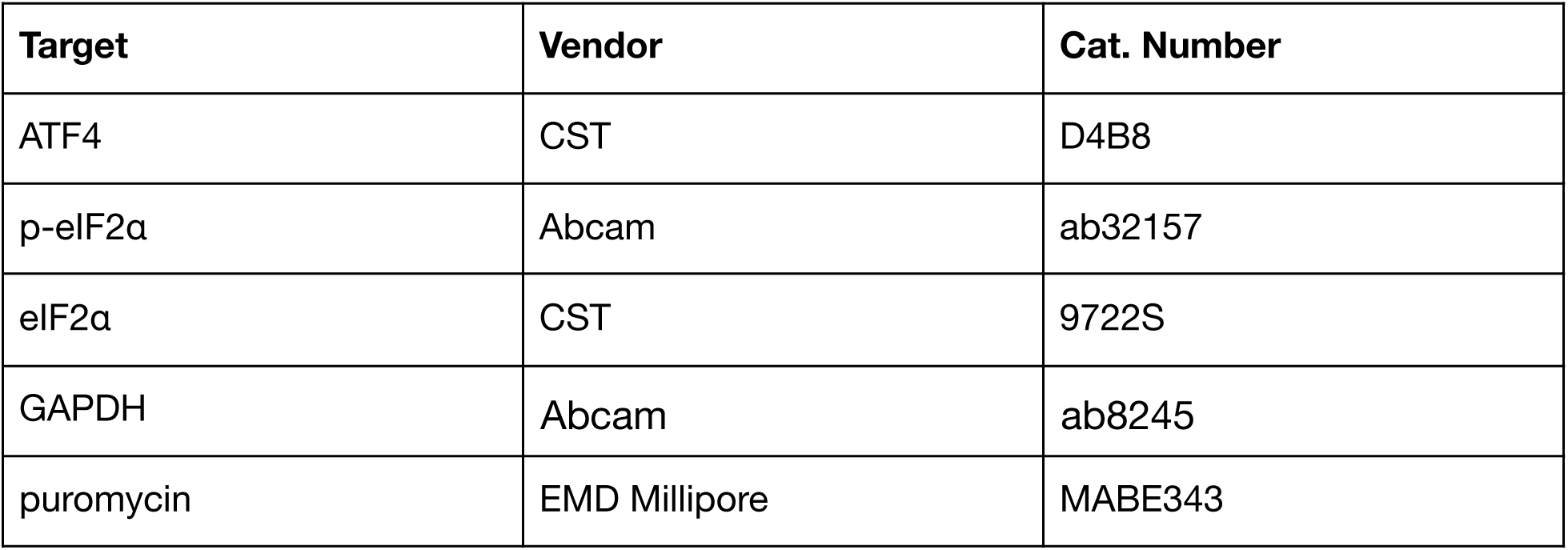

| Target | Vendor | Cat. Number |
| --- | --- | --- |
| ATF4 | CST | D4B8 |
| p-eIF2α | Abcam | ab32157 |
| eIF2α | CST | 9722S |
| GAPDH | Abcam | ab8245 |
| puromycin | EMD Millipore | MABE343 |

#### c) Puromycin incorporation assay

H4 FKBP-PKR cells were grown in 6-well plates at 70% confluency, and the AP20187 homodimerizer was added starting 16 hours later for the indicated time points. 1µg/ml of puromycin was spiked into the different wells for 30 min at 37°C and 5% CO2. Cells were collected and processed as described for immunoblotting.

### Model implementation

We implemented the model using the open-source Python package PySB (Lopez et al., 2013), which is a framework for building rule-based mathematical models of biochemical systems. The primary model consists of 28 species and 56 free parameters, whereas the simplified model consists of 21 species and 35 free parameters. All the reactions considered in the primary model are listed in Table S1, including their forward and reverse reaction rates. The initial values of the model components were set based on the proteomics data. Note that the control cells didn’t have the FKBP-PKR tool in our experiment. Therefore, we couldn’t use the wild-type cell states as the initial state. Instead, we used 0-hour measurements of drugged cells as the initial values, which were collected right before the drug administration from cells with the FKBP-PKR tool.

### Model calibration

We used the open-source Python package simplePSO to calibrate the model’s free parameters (Pino, 2019), which implements the meta-heuristic particle swarm optimization algorithm (Gad, 2022). We arbitrarily initialized model parameters, considering larger/smaller values for binding/unbinding reactions. Then, for each free parameter *p* except the p-eIF2α to total eIF2α ratio threshold (which is between 0 and 1), we scanned [*log*(*p*) − 4, *log*(*p*) + 4], while minimizing the total mean squared error between simulated and experimental ATF4, GADD34, CHOP, and DR5, i.e.

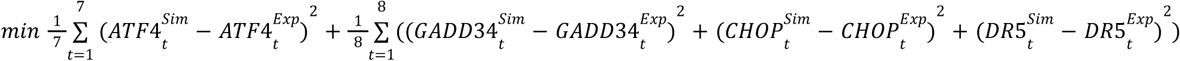

Note that we excluded the ATF4 measurement at the 8th hour because ATF4 was 0 in all samples. Based on our previous experiments (Zappa et al., 2022), we don’t expect ATF4 to be detected in the 6th-hour samples, degrade in the 8th-hour samples, and peak again in the 16th-hour samples. It is likely that the mass spectrometer did not detect ATF4 at the 8th hour. That’s why the summation for ATF4 runs from *t* = 1 to 7 to indicate the 7 time points. Additionally, we min-max normalized the experimental data (and simulation output accordingly) to eliminate the effects of differences in scale of protein abundances on efficiency of search space scanning, and to accelerate convergence to a solution while solving the optimization problem. We ran the optimization several times using different starting points in the high-dimensional parameter space to make sure the search space was well-scanned. See supplementary files for the estimated parameter values of the best-fitting model.

### Sensitivity analysis

The model sensitivity analyses were done using the Sobol method (Sobol, 2001), which is a variance-based sensitivity analysis that decomposes the variance at the output of the model to the changes in its input(s). We used the open-source Python package SALib (Herman and Usher, 2017), in which Sobol and some other methods are implemented. Overall, the Sobol method outputs three values: S1, S2, and ST. S1 is the first-order sensitivity index, which measures the contribution to the output variance of the change in the individual initial value. S2 is the second-order sensitivity index, which measures the contribution to the output variance of the change in two initial values. Lastly, ST is the total-effect index, which measures the contribution to the output variance caused by an individual initial value, including its first-order effects (the input varying alone) and all higher-order interactions (the input varying together with other input combinations).

To analyze model sensitivity to the initial values, we considered a 30% deviation from the experimental 0-hour values and quantified how the variance at the output fitness to experimental data (i.e., the variance in MSE value) changes after 24 hours of simulation. To elaborate, for each species *S*, we generated 65K samples via Saltelli sampling (Saltelli, 2002) in the range [0. 7 * [*S*], 1. 3 * [*S*]] if *S* > 0, and [0, 1*e*3] otherwise. Next, we simulated the pathway for each of these samples and computed the MSE value between the output and its experimental value. Then, based on the variance in the MSE values, the sensitivity value of the species *S* is computed using the Sobol method. Similarly, to assess the model’s sensitivity to the free parameters, we generated 65K samples in the range [0. 5 * *p*, 2 * *p*] for each parameter *p*, ran the simulation for each sample, and then computed the MSE value between the output and its experimental value. Lastly, we applied the Sobol method to the MSE values. See supplementary spreadsheets for S1, S2, and ST values of both initial value and parameter sensitivities.

## Acknowledgments

We would like to thank Lucas Reineke, Advait Subramanian, Morgane Boone, William Poole, and Chaitra Agrahar for insightful discussion and critical feedback on this work.

## Code and data availability

The model and the downstream analyses were implemented in Python, designed to be used with Jupyter Notebooks. Source files, experimental data, and step-by-step instructions on how to run the model can be found in our GitHub repository: https://github.com/altoslabs/isrmm

## Author contributions

D.A-A and C.F.L supervised the research. M.O. developed the methods, performed the simulations and computations. F.Z. designed experiments. F.Z., and K.K. performed experiments. D.I. performed mass spectrometry. S.T., F.Z., and M.O. analyzed the data. M.O., F.Z., D.I., S.T., C.F.L., and D.A-A wrote the manuscript.

## References

Abdulkarim, B., Nicolino, M., Igoillo-Esteve, M., Daures, M., Romero, S., Philippi, A., Senée, V., Lopes, M., et al. (2015). A missense mutation in PPP1R15B causes a syndrome including diabetes, short stature, and microcephaly. Diabetes 64(11):3951–3962. 10.2337/db15-0477

Adamson, B., Norman, T.M., Jost, M., Cho, M.Y., Nuñez, J.K., Chen, Y., Villalta, J.E., Gilbert, L.A., et al. (2016). A multiplexed single-cell CRISPR screening platform enables systematic dissection of the unfolded protein response. Cell 167(7):1867–1882.e21. 10.1016/j.cell.2016.11.048

Albeck, J.G., Burke, J.M., Spencer, S.L., Lauffenburger, D.A., and Sorger, P.K. (2008). Modeling a snap-action, variable-delay switch controlling extrinsic cell death. PLoS Biol. 6(12): 2831–2852. 10.1371/journal.pbio.0060299

Andreev, D.E., O’Connor, P.B., Fahey, C., Kenny, E.M., Terenin, I.M., Dmitriev, S.E., Cormican, P., Morris, D.W., Shatsky, I.N., and Baranov, P.V. (2015). Translation of 5’ leaders is pervasive in genes resistant to eIF2 repression. eLife 4, e03971. 10.7554/eLife.03971

Averous, J., Bruhat, A., Jousse, C., Carraro, V., Thiel, G., and Fafournoux, P. (2004). Induction of CHOP expression by amino acid limitation requires both ATF4 expression and ATF2 phosphorylation. J. Biol. Chem. 279(7): 5288–5297. 10.1074/jbc.M311862200

Axten, J.M., Romeril, S.P., Shu, A., Ralph, J., Medina, J.R., Feng, Y., Li, W.H., Grant, S.W., et al. (2013). Discovery of GSK2656157: An optimized PERK inhibitor selected for preclinical development. ACS Med. Chem. Lett. 4(10):964–968. 10.1021/ml400228e

Batjargal, T., Zappa, F., Grant, R.J., Piscopio, R.A., Chialastri, A., Dey, S.S., Acosta-Alvear, D., and Wilson, M.Z. (2023). Optogenetic control of the integrated stress response reveals proportional encoding and the stress memory landscape. Cell Syst. 14(7):551–562.e5. 10.1016/j.cels.2023.06.001

Boone, M., and Zappa, F. (2023). Signaling plasticity in the integrated stress response. Front. Cell Dev. Biol. 11:1271141. 10.3389/fcell.2023.1271141

Brar, K.K., Hughes, D.T., Morris, J.L., Subramanian, K., Krishna, S., Gao, F., Rieder, L.S., Uhrig, S., et al. (2024). PERK-ATAD3A interaction provides a subcellular safe haven for protein synthesis during ER stress. Science 385(6712), eadp7114. 10.1126/science.adp7114

Brito Querido, J., Díaz-López, I., and Ramakrishnan, V. (2024). The molecular basis of translation initiation and its regulation in eukaryotes. Nat Rev Mol Cell Biol. 25(3):168–186. 10.1038/s41580-023-00624-9

Burkart, S.S., Schweinoch, D., Frankish, J., Sparn, C., Wüst, S., Urban, C., Merlo, M., Magalhães, V.G., Piras, A., Pichlmair, A., Willemsen, J., Kaderali, L., and Binder, M. (2023). High-resolution kinetic characterization of the RIG-I-signaling pathway and the antiviral response. Life Sci. Alliance 6(10):e202302059. 10.26508/lsa.202302059

Chakrabarty, Y., Yang, Z., Chen, H., and Chan, D.C. (2024). The HRI branch of the integrated stress response selectively triggers mitophagy. Mol. Cell 84(6):1090–1100.e6. 10.1016/j.molcel.2024.01.016

Clackson, T., Yang, W., Rozamus, L.W., Hatada, M., Amara, J.F., Rollins, C.T., Stevenson, L.F., Magari, S.R. et al. (1998). Re-designing an FKBP-ligand interface to generate chemical dimerizers with novel specificity. Proc. Natl. Acad. Sci. USA. 95:10437–10442. 10.1073/pnas.95.18.10437

Clemens M.J. (1997). PKR–a protein kinase regulated by double-stranded RNA. Int. J. Biochem. & Cell Biol 29(7):945–949. 10.1016/s1357-2725(96)00169-0

Clemens, M.J., Pain, V.M., Wong, S.T., and Henshaw, E.C. (1982). Phosphorylation inhibits guanine nucleotide exchange on eukaryotic initiation factor 2. Nature 296(5852):93–95. 10.1038/296093a0

Cole J.L. (2007). Activation of PKR: an open and shut case?. Trends Biochem Sci. 32(2):57–62. 10.1016/j.tibs.2006.12.003

Connor, J.H., Weiser, D.C., Li, S., Hallenbeck, J.M., and Shenolikar, S. (2001). Growth arrest and DNA damage-inducible protein GADD34 assembles a novel signaling complex containing protein phosphatase 1 and inhibitor 1. Mol. Cell. Biol. 21(20):6841–6850. 10.1128/MCB.21.20.6841-6850.2001

Costa-Mattioli, M., and Walter, P. (2020). The integrated stress response: From mechanism to disease. Science. 368:eaat5314. 10.1126/science.aat5314

Dar, A.C., Dever, T.E., and Sicheri, F. (2005). Higher-order substrate recognition of eIF2alpha by the RNA-dependent protein kinase PKR. Cell 122(6):887–900. 10.1016/j.cell.2005.06.044

Donovan, J., Whitney, G., Rath, S., and Korennykh, A. (2015). Structural mechanism of sensing long dsRNA via a noncatalytic domain in human oligoadenylate synthetase 3. Proc. Natl. Acad. Sci. USA. 112(13):3949–3954. 10.1073/pnas.1419409112

Eling, N., Morgan, M.D., and Marioni, J.C. (2019). Challenges in measuring and understanding biological noise. Nat. Rev. Genet. 20:536–548. 10.1038/s41576-019-0130-6

Emadi, A., Ozen, M., and Abdi, A. (2022).A hybrid model to study how late long-term potentiation is affected by faulty molecules in an intraneuronal signaling network regulating transcription factor CREB. Integr. Biol. 14(5):111–125. 10.1093/intbio/zyac011

Feng, G.S., Chong, K., Kumar, A., and Williams, B.R. (1992). Identification of double-stranded RNA-binding domains in the interferon-induced double-stranded RNA-activated p68 kinase. Proc. Natl. Acad. Sci. USA. 89(12): 5447–5451. 10.1073/pnas.89.12.5447

Gad, A.G. (2022). Particle swarm optimization algorithm and its applications: A systematic review. Arch. Computat. Methods. Eng. 29:2531–2561. 10.1007/s11831-021-09694-4

Gomez, E., Mohammad, S.S. and Pavitt, G.D. (2002). Characterization of the minimal catalytic domain within eIF2B: the guanine-nucleotide exchange factor for translation initiation. EMBO J. 21(19):5292–5301. 10.1093/emboj/cdf515

Gordiyenko, Y., Llácer, J.L., and Ramakrishnan, V. (2019). Structural basis for the inhibition of translation through eIF2α phosphorylation. Nat. Commun. 10:2640. 10.1038/s41467-019-10606-1

Gordiyenko, Y., Schmidt, C., Jennings, M.D., Matak-Vinkovic, D., Pavitt, G.D., and Robinson, C.V. (2014). eIF2B is a decameric guanine nucleotide exchange factor with a γ2ε2 tetrameric core. Nat. Commun. 5, 3902. 10.1038/ncomms4902

Guilbert, M., Anquez, F., Pruvost, A., Thommen, Q., and Courtade, E. (2020). Protein level variability determines phenotypic heterogeneity in proteotoxic stress response. FEBS J. 287(24):5345–5361. 10.1111/febs.15297

Harding, H.P., Novoa, I., Zhang, Y., Zeng, H., Wek, R., Schapira, M., and Ron, D. (2000). Regulated translation initiation controls stress-induced gene expression in mammalian cells. Mol Cell 6:1099–1108. 10.1016/S1097-2765(00)00108-8

Herman, J., and Usher, W. (2017). SALib: An open-source Python library for sensitivity analysis. Journal of Open Source Software, 2(9). 10.21105/joss.00097

Hinnebusch, A.G., Ivanov, I.P., and Sonenberg, N. (2016). Translational control by 5′-untranslated regions of eukaryotic mRNAs. Science 352:1413–1416. 10.1126/science.aad9868

Holcik, M., and Sonenberg, N. (2005). Translational control in stress and apoptosis. Nat. Rev. Mol. Cell Biol. 6, 318–327. 10.1038/nrm1618

Hotamisligil, G.S., and Davis, R.J. (2016). Cell Signaling and Stress Responses. Cold Spring Harb Perspect Biol. 8(10):a006072. 10.1101/cshperspect.a006072

Hu, H., Tian, M., Ding, C., and Yu, S. (2019). The C/EBP homologous protein (CHOP) transcription factor functions in endoplasmic reticulum stress-induced apoptosis and microbial infection. Front Immunol. 9:3083. 10.3389/fimmu.2018.03083

Je, H.S., Lu, Y., Yang, F., Nagappan, G., Zhou, J., Jiang, Z., Nakazawa, K., and Lu, B. (2009). Chemically inducible inactivation of protein synthesis in genetically targeted neurons. J. Neurosci. 29(21):6761–6766. 10.1523/JNEUROSCI.1280-09.2009

Jiang, Z., Belforte, J.E., Lu, Y., Yabe, Y., Pickel, J., Smith, C.B., Je, H.S., Lu, B., and Nakazawa, K. (2010). eIF2alpha Phosphorylation-dependent translation in CA1 pyramidal cells impairs hippocampal memory consolidation without affecting general translation. J. Neurosci. 30(7):2582–2594. 10.1523/JNEUROSCI.3971-09.2010

Jousse, C., Oyadomari, S., Novoa, I., Lu, P., Zhang, Y., Harding, H.P., and Ron, D. (2003). Inhibition of a constitutive translation initiation factor 2alpha phosphatase, CReP, promotes survival of stressed cells. J. Cell Biol. 163(4):767–775. 10.1083/jcb.200308075

Kapp, L.D. and Lorsch, J.R. (2004). GTP-dependent recognition of the methionine moiety on initiator tRNA by translation factor eIF2. J. Mol. Biol. 335(4):923–936. 10.1016/j.jmb.2003.11.025

Kojima, E., Takeuchi, A., Haneda, M., Yagi, A., Hasegawa, T., Yamaki, K., Takeda, K., Akira, S., et al. (2003). The function of GADD34 is a recovery from a shutoff of protein synthesis induced by ER stress: Elucidation by GADD34-deficient mice. FASEB J. 17:1573–1575. 10.1096/fj.02-1184fje

Klein, P., Kallenberger, S.M., Roth, H., Roth, K., Ly-Hartig, T.B.N., Magg, V., Aleš, J., Talemi, S.R. et al. (2022). Temporal control of the integrated stress response by a stochastic molecular switch. Sci Adv. 8(12):eabk2022. 10.1126/sciadv.abk2022

Kourtis, N., and Tavernarakis, N. (2011). Cellular stress response pathways and ageing: intricate molecular relationships. EMBO J. 30(13):2520–2531. 10.1038/emboj.2011.162

Krishnamoorthy, T., Pavitt, G.D., Zhang, F., Dever, T.E., and Hinnebusch, A.G. (2001). Tight binding of the phosphorylated α subunit of initiation factor 2 (eIF2α) to the regulatory subunits of guanine nucleotide exchange factor eIF2B is required for inhibition of translation initiation. Mol. Cell Biol. 21(15):5018–5030. 10.1128/MCB.21.15.5018-5030.2001

Lassot, I., Ségéral, E., Berlioz-Torrent, C., Durand, H., Groussin, L., Hai, T., Benarous, R., and Margottin-Goguet, F. (2001). ATF4 degradation relies on a phosphorylation-dependent interaction with the SCF(betaTrCP) ubiquitin ligase. Mol. Cell Biol. 21(6):2192–2202. 10.1128/MCB.21.6.2192-2202.2001

Li, S., Peters, G.A., Ding, K., Zhang, X., Qin, J., and Sen, G.C. (2006). Molecular basis for PKR activation by PACT or dsRNA. Proc. Natl. Acad. Sci. USA. 103(26):10005–10010. 10.1073/pnas.0602317103

Linden, N.J., Kramer, B., and Rangamani, P. (2022). Bayesian parameter estimation for dynamical models in systems biology. PLoS Comput. Biol. 18(10):e1010651. 10.1371/journal.pcbi.1010651

Lopez, C.F., Muhlich, J.L., Bachman, J.A., and Sorger, P.K. (2013). Programming biological models in Python using PySB. Mol Syst Biol. 9:646. 10.1038/msb.2013.1

Lu, P.D., Harding, H.P., and Ron, D. (2004). Translation reinitiation at alternative open reading frames regulates gene expression in an integrated stress response. J Cell Biol. 167(1):27–33. 10.1083/jcb.200408003

Lu, P.D., Jousse, C., Marciniak, S.J., Zhang, Y., Novoa, I., Scheuner, D., Kaufman, R.J., Ron, D., and Harding, H.P. (2004). Cytoprotection by pre-emptive conditional phosphorylation of translation initiation factor 2. EMBO J. 23(1):169–179. 10.1038/sj.emboj.7600030

Lu, M., Lawrence, D.A., Marsters, S., Acosta-Alvear, D., Kimmig, P., Mendez, A.S., Paton, A.W., Paton, J.C., Walter, P., and Ashkenazi, A. (2014). Opposing unfolded-protein-response signals converge on death receptor 5 to control apoptosis. Science 345(6192):98–101. 10.1126/science.1254312

Ma, Y., and Hendershot, L.M. (2003). Delineation of a negative feedback regulatory loop that controls protein translation during endoplasmic reticulum stress. J. Biol. Chem. 278(37):34864–34873. 10.1074/jbc.M301107200

Mayo, C.B., Erlandsen, H., Mouser, D.J., Feinstein, A.G., Robinson, V.L., May, E.R., and Cole, J.L. (2019). Structural basis of protein kinase R autophosphorylation. Biochemistry 58(27):2967–2977. 10.1021/acs.biochem.9b00161

Nanduri, S., Carpick, B.W., Yang, Y., Williams, B.R., and Qin, J. (1998). Structure of the double-stranded RNA-binding domain of the protein kinase PKR reveals the molecular basis of its dsRNA-mediated activation. EMBO J 17(18):5458–5465. 10.1093/emboj/17.18.5458

Novoa, I., Zeng, H., Harding, H.P., and Ron, D. (2001). Feedback inhibition of the unfolded protein response by GADD34-mediated dephosphorylation of eIF2alpha. J Cell Biol. 153(5), 1011–1022. 10.1083/jcb.153.5.1011

Ortega, O.O., Ozen, M., Wilson, B.A., Pino, J.C., Irvin, M.W., Ildefonso, G.V., Garbett, S.P., and Lopez, C.F. (2024). Signal execution modes emerge in biochemical reaction networks calibrated to experimental data. iScience 27(6):109989. 10.1016/j.isci.2024.109989

Ozen, M., Lipniacki, T., Levchenko, A., Emamian, E.S., and Abdi, A. (2020). Modeling and measurement of signaling outcomes affecting decision making in noisy intracellular networks using machine learning methods. Integr. Biol. 12(5):122–138. 10.1093/intbio/zyaa009

Pakos-Zebrucka, K., Koryga, I., Mnich, K., Ljujic, M., Samali, A., and Gorman, A.M. (2016). The integrated stress response. EMBO Rep. 17(10):1374–1395. 10.15252/embr.201642195

Pino, J. (2019). simplePSO: Simple interface to optimize models using Particle Swarm Optimization [Software]. GitHub. https://github.com/LoLab-MSM/simplePSO

Rojas, M., Vasconcelos, G., and Dever, T.E. (2015). An eIF2α-binding motif in protein phosphatase 1 subunit GADD34 and its viral orthologs is required to promote dephosphorylation of eIF2α. Proc. Natl. Acad. Sci. USA. 112(27):E3466–E3475. 10.1073/pnas.1501557112

Rowlands, A.G., Panniers, R., and Henshaw, E.C. (1988). The catalytic mechanism of guanine nucleotide exchange factor action and competitive inhibition by phosphorylated eukaryotic initiation factor 2. J. Biol. Chem. 263(12):5526–5533.

Saltelli, A. (2002). Making best use of model evaluations to compute sensitivity indices. Comp. Phys. Comm. 145(2):280–297. 10.1016/S0010-4655(02)00280-1

Sevrin, T., Imoto, H., Robertson, S., Rauch, N., Dyn’ko, U., Koubova, K., Wynne, K., Kolch, W., Rukhlenko, O.S., and Kholodenko, B.N. (2024). Cell-specific models reveal conformation-specific RAF inhibitor combinations that synergistically inhibit ERK signaling in pancreatic cancer cells. Cell Reports, 43(9):114710. 10.1016/j.celrep.2024.114710

Shockley, E.M., Rouzer, C.A., Marnett, L.J., Deeds, E.J., and Lopez, C.F. (2019). Signal integration and information transfer in an allosterically regulated network. npj Syst. Biol. Appl. 5, 23. 10.1038/s41540-019-0100-9.

Sidrauski, C., McGeachy, A.M., Ingolia, N.T., and Walter, P. (2015). The small molecule ISRIB reverses the effects of eIF2α phosphorylation on translation and stress granule assembly. eLife 4, e05033. 10.7554/eLife.05033

Sobol, I.M. (2001). Global sensitivity indices for nonlinear mathematical models and their Monte Carlo estimates. Math. and Comp. in Sim. 55(1-3):271–280. 10.1016/S0378-4754(00)00270-6

Szaruga, M., Janssen, D.A., de Miguel, C. et al. Activation of the integrated stress response by inhibitors of its kinases. Nat. Commun. 14, 5535 (2023). 10.1038/s41467-023-40823-8

Tang, C.P., Clark, O., Ferrarone, J.R., Campos, C., Lalani, A.S., Chodera, J.D., Intlekofer, A.M., Elemento, O., and Mellinghoff, I.K. (2022). GCN2 kinase activation by ATP-competitive kinase inhibitors. Nat. Chem. Biol. 18(2):207–215. 10.1038/s41589-021-00947-8

Tian, X., Zhang, S., Zhou, L., Seyhan, A.A., Hernandez Borrero, L., Zhang, Y., and El-Deiry, W.S. (2021). Targeting the integrated stress response in cancer therapy. Front Pharmacol. 12:747837. 10.3389/fphar.2021.747837

Torkenczy, K., Reineke, L.C., Dooling, S.W., Henderson, B., Itzhak, D.N., et al. (2025). Mapping the ISR landscape in cognitive disorders via single-cell multi-omics. bioRxiv 2025.02.28.640905. 10.1101/2025.02.28.640905

UniProt Consortium (2025). UniProt: the Universal Protein Knowledgebase in 2025. Nucleic Acids Res. 53(D1):D609–D617. 10.1093/nar/gkae1010

Wiśniewski, J.R., Hein, M.Y., Cox, J., and Mann, M. (2014). A "proteomic ruler" for protein copy number and concentration estimation without spike-in standards. Mol. Cell. Proteomics 13(12):3497–3506. 10.1074/mcp.M113.037309

Wong, F., Li, A., Omori, S., Lach, R.S., Nunez, J., Ren, Y., Brown, S.P., Singhal, V., Lyda, B.R., Batjargal, T., Dickson, E., Reyes, J.R.R., Vargas, J.M.U., Wahane, S., Kim, H., Collins, J.J., and Wilson, M.Z. (2025). Optogenetics-enabled discovery of integrated stress response modulators. Cell 188:1–18. 10.1016/j.cell.2025.06.024

Wong, Y.L., LeBon, L., Basso, A.M., Kohlhaas, K.L., Nikkel, A.L., Robb, H.M., Donnelly-Roberts, D.L., Prakash, J., et al. (2019). eIF2B activator prevents neurological defects caused by a chronic integrated stress response. eLife 8, e42940. 10.7554/eLife.42940

Wolozin, B. and Ivanov, P. (2019). Stress granules and neurodegeneration. Nat Rev Neurosci. 20(11):649–666. 10.1038/s41583-019-0222-5

Xu, X., Krumm, C., So, J.S., Bare, C.J., Holman, C., Gromada, J., Cohen, D.E., & Lee, A. H. (2018). Preemptive activation of the integrated stress response protects mice from diet-Induced obesity and insulin resistance by fibroblast growth factor 21 Induction. Hepatology 68(6):2167–2181. 10.1002/hep.30060

Yamaguchi, H., and Wang, H.G. (2004). CHOP is involved in endoplasmic reticulum stress-induced apoptosis by enhancing DR5 expression in human carcinoma cells. J. Biol. Chem. 279(44):45495–45502. 10.1074/jbc.M406933200

Yu, Q., Qu, K., and Modis, Y. (2018). Cryo-EM structures of MDA5-dsRNA filaments at different stages of ATP hydrolysis. Mol. Cell 72(6):999–1012.e6. 10.1016/j.molcel.2018.10.012

Zappa, F., Muniozguren, N.L., Conrad, J.E., and Acosta-Alvear, D. (2025). The integrated stress response engages a cell-autonomous, ligand-independent, DR5-driven apoptosis switch. Cell Death Dis. 16:101. 10.1038/s41419-025-07403-8

Zappa, F., Muniozguren, N.L., Wilson, M.Z., Costello, M.S., Ponce-Rojas, J.C., and Acosta-Alvear, D. (2022). Signaling by the integrated stress response kinase PKR is fine-tuned by dynamic clustering. J Cell Biol. 221(7):e202111100. 10.1083/jcb.202111100

Zhang, G., Wang, X., Rothermel, B.A., Lavandero, S., and Wang, Z.V. (2022). The integrated stress response in ischemic diseases. Cell Death Differ. 29(4):750–757. 10.1038/s41418-021-00889-7

