## Supplementary material for "Intrinsic Molecular Timers and a Biphasic Amplitude Limit Regulate the Integrated Stress Response": Ozen et al_2025_Supplementary information.pdf

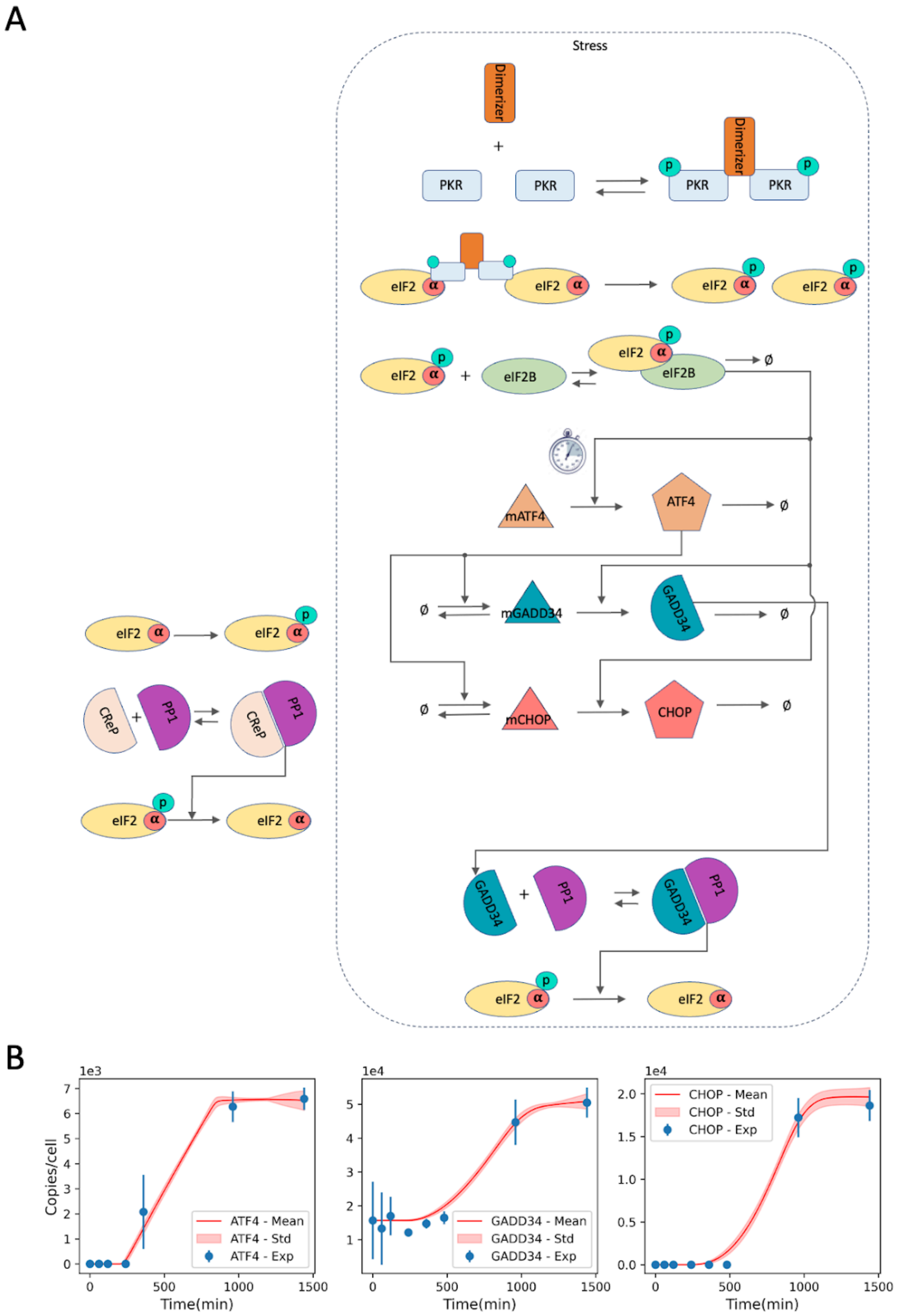


**Figure S1: Predicting switch p-eIF2α to total eIF2α ratio threshold.** (A) The simplified ISR model. (B) The goodness of fit of the model for the observed 97 parameter sets via multi-start optimization.

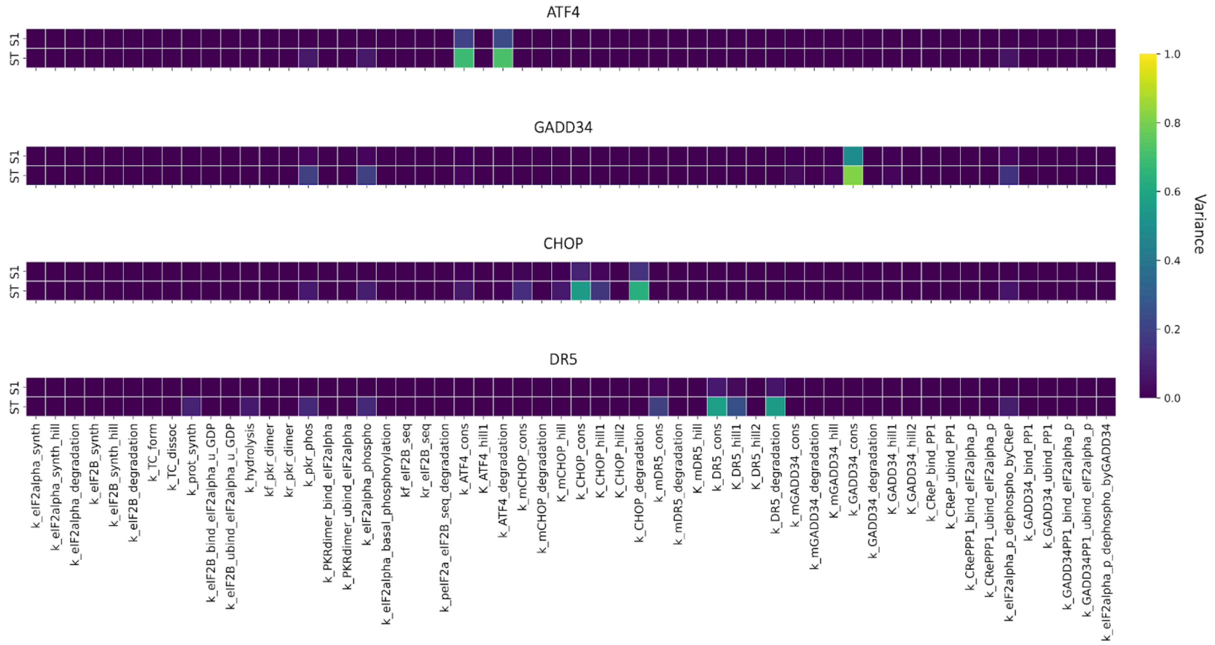


**Figure S2: Model sensitivity analysis based on its free parameters.** The model's sensitivity, estimated using the free parameters of the system of ordinary differential equations (Table S1) representing the entire network of interactions between core ISR components (Figure 1), was computed for ATF4, CHOP, GADD34, and DR5 independently. In these calculations, these four proteins represent model outputs, while the free parameters were considered inputs. The sensitivity values were computed based on the goodness of fit of the experimental data to the model's output (i.e., variance in the MSE between output and the experimental data, as indicated by the colorbar). A high variance indicates that the model is highly sensitive to a particular input. S1 measures the contribution to the output variance of the change in an individual free parameter. ST measures the contribution to the output variance caused by an individual free parameter, including both its first-order effects (variance of the free parameter) and all higher-order interactions (covariance of the free parameter alongside combinations of other free parameters).

**Table S1. Reactions and reaction rates in the primary ISR model.** ":" indicates complexation. "⇌" indicates reversible reactions, whereas "⟶" indicates forward reactions. For modeling purposes, we considered eIF2α binding with GTP/GDP as one of the eIF2α states. Therefore, eIF2α(GTP) and eIF2α(GDP) indicate such bindings accordingly. The simplified ISR model presented in Figure S2 excludes the colored reactions.

| Reaction | Forward rate | Reverse rate |
|---|---|---|
| ∅ ⇌ u-eIF2α(GDP) | $k_1 \times \frac{[u-eIF2\alpha(GTP):Met-tRNAi]}{K_1+[u-eIF2\alpha(GTP):Met-tRNAi]}$ | $k_2 \times$ [u-eIF2α(GDP)] |
| ∅ ⇌ eIF2B | $k_3 \times \frac{[u-eIF2\alpha(GTP):Met-tRNAi]}{K_2+[u-eIF2\alpha(GTP):Met-tRNAi]}$ | $k_4 \times$ [eIF2B] |
| u-eIF2α(GTP) + Met-tRNAi ⇌ u-eIF2α(GTP):Met-tRNAi | $k_5 \times$ [u-eIF2α(GTP)] × [Met-tRNAi] | $k_6 \times$ [u-eIF2α(GTP):Met-tRNAi] |
| u-eIF2α(GTP):Met-tRNAi ⟶ u-eIF2α(GDP) + Protein | $k_7 \times$ [u-eIF2α(GTP):Met-tRNAi] | — |
| u-eIF2α(GDP) + eIF2B ⇌ u-eIF2α(GDP):eIF2B | $k_8 \times$ [u-eIF2α(GDP)] × [eIF2B] | $k_9 \times$ [u-eIF2α(GDP):eIF2B] |
| u-eIF2α(GDP):eIF2B ⟶ u-eIF2α(GTP) + eIF2B | $k_{10} \times$ [u-eIF2α(GDP):eIF2B] | — |
| u-eIF2α(GDP) ⟶ p-eIF2α | $k_{11} \times$ [u-eIF2α(GDP)] | — |
| CReP + PP1 ⇌ CReP:PP1 | $k_{12} \times$ [CReP] × [PP1] | $k_{13} \times$ [CReP:PP1] |
| CReP:PP1 + p-eIF2α ⇌ CReP:PP1:p-eIF2α | $k_{14} \times$ [CReP:PP1] × [p-eIF2α] | $k_{15} \times$ [CReP:PP1:p-eIF2α] |
| CReP:PP1:p-eIF2α ⟶ CReP:PP1 + u-eIF2α(GDP) | $k_{16} \times$ [CReP:PP1:p-eIF2α] | — |
| u-PKR + Dimerizer + u-PKR ⇌ u-PKR:Dimerizer:u-PKR | $k_{17} \times$ [u-PKR]$^2$ × [Dimerizer] | $k_{18} \times$ [u-PKR:Dimerizer:u-PKR] |

| | | |
|---|---|---|
| u-PKR:Dimerizer:u-PKR ⟶ p-PKR:Dimerizer:p-PKR | $k_{19}$ × [u-PKR:Dimerizer:u-PKR] | — |
| p-PKR:Dimerizer:p-PKR + 2u-eIF2α(GDP) ⇌ p-PKR:Dimerizer:p-PKR:2u-eIF2α(GDP) | $k_{20} \times [u - eIF2\alpha(GDP)]^2$ × [p-PKR:Dimerizer:p-PKR] | $k_{21}$ × [p-PKR:Dimerizer:p-PKR:2u-eIF2α(GDP)] |
| p-PKR:Dimerizer:p-PKR:2u-eIF2α(GDP) ⟶ p-PKR:Dimerizer:p-PKR + 2p-eIF2α | $k_{22}$ × [p-PKR:Dimerizer:p-PKR:2u-eIF2α (GDP)] | — |
| p-eIF2α + eIF2B ⇌ p-eIF2α:eIF2B | $k_{23}$ × [p-eIF2α] × [eIF2B] | $k_{24}$ × [p-eIF2α:eIF2B] |
| p-eIF2α:eIF2B ⟶ ∅ | $k_{25}$ × [p-eIF2α:eIF2B] | — |
| ∅ ⇌ ATF4 | $k_{26} \times 1(\frac{[p-eIF2\alpha]}{[eIF2\alpha]} > 0.3)) \times \frac{[p-eIF2\alpha:eIF2B]}{K_3+[p-eIF2\alpha:eIF2B]}$ | $k_{27}$ × [ATF4] |
| ∅ ⇌ mGADD34 | $k_{28} \times \frac{[ATF4]}{K_4+[ATF4]}$ | $k_{29}$ × [mGADD34] |
| ∅ ⇌ GADD34 | $k_{30} \times \frac{[mGADD34]}{K_5+[mGADD34]} \times \frac{[p-eIF2\alpha:eIF2B]}{K_6+[p-eIF2\alpha:eIF2B]}$ | $k_{31}$ × [GADD34] |
| ∅ ⇌ mCHOP | $k_{32} \times \frac{[ATF4]}{K_7+[ATF4]}$ | $k_{33}$ × [mCHOP] |
| ∅ ⇌ CHOP | $k_{34} \times \frac{[mCHOP]}{K_8+[mCHOP]}$ | $k_{35}$ × [CHOP] |

| | | |
|---|---|---|
| | $\times \frac{[p-eIF2\alpha:eIF2B]}{K_9+[p-eIF2\alpha:eIF2B]}$ | |
| Ø ⇌ mDR5 | $k_{36} \times \frac{[CHOP]}{K_{10}+[CHOP]}$ | $k_{37}$ × [mDR5] |
| Ø ⇌ DR5 | $k_{38} \times \frac{[mDR5]}{K_{11}+[mDR5]}$ $\times \frac{[u-eIF2\alpha(GTP):Met-tRNAi]}{K_{12}+[u-eIF2\alpha(GTP):Met-tRNAi]}$ | $k_{39}$ × [DR5] |
| GADD34 + PP1 ⇌ GADD34:PP1 | $k_{40}$ × [GADD34] × [PP1] | $k_{41}$ × [GADD34:PP1] |
| GADD34:PP1 + p-eIF2α ⇌ GADD34:PP1:p-eIF2α | $k_{42}$ × [GADD34:PP1] × [p-eIF2α] | $k_{43}$ × [GADD34:PP1:p-eIF2α] |
| GADD34:PP1:p-eIF2α ⟶ GADD34:PP1 + u-eIF2α(GDP) | $k_{44}$ × [GADD34:PP1:p-eIF2α] | — |
